# Extracting biological structure and heterogeneity from the nano to the macro scale

**DOI:** 10.1101/2025.08.06.668903

**Authors:** Edward Rosten, Daniel Stedman, Lee-Ya Chu, Kokila Wickramanayake, George Littlejohn, Katherine Baxter, Gail McConnell, Siân Culley, Queelim Ch’ng, Christophe Leterrier, Mark Bates, Maxim Molodtsov, Susan Cox

**Affiliations:** Randall Centre for Cell and Molecular Biophysics, King’s College London, London, United Kingdom; The Francis Crick Institute, London, United Kingdom; School of Biological and Marine Sciences, University of Plymouth, Plymouth, United Kingdom; Strathclyde Institute of Pharmacy and Biomedical Sciences, University of Strathclyde, Glasgow, United Kingdom; Centre for Developmental Neurobiology, King’s College London, London, United Kingdom; NeuroCyto, Aix Marseille Université, Marseille, France; Department of Optical Nanoscopy, Institute for Nanophotonics, Göttingen, Germany; Department of Physics and Astronomy, University College London, London, United Kingdom

## Abstract

Fluorescence microscopy is an essential tool in biology. It has revealed great variability at multiple scales, in macromolecular complexes, cells, and organisms. Understanding this variability will reveal the mechanisms by which genetically or biochemically identical systems adopt different biological states. Achieving this requires the ability to extract both the underlying biological structure and how it varies across the population. Currently the field lacks general techniques to deal with arbitrary structures and different types of variability. Here we present SQUASSH, a new convolutional neural network-based approach to freely fit structural models to fluorescence microscopy data that simultaneously quantifies variability to reveal correlations, dynamics, and systematic distortions. SQUASSH is highly versatile: it accommodates diverse imaging modalities at length scales from nm to mm. This approach opens up applications such as imaging nanoscale macromolecular structures, revealing patterns in shape changes from organelle to tissue scale, and characterizing systems biology of dynamical processes.

## 1 Introduction

In biology, observation enables discovery, and so throughout its history biological discovery has been driven by technical innovations in imaging, particularly microscopy. Fluorescence microscopy allows specific molecules to be labelled and thus imaged, with recent developments having pushed the achievable resolution to 1 nm ^1^. It is now possible to acquire large volumes of data, either observations of many instances of the same type of structure (sub-cellular structure, entire cell, or organism) or of the same structure observed over time. When the structure is thought to be identical in each observation information from all the observations can be combined into a single, more accurate structure using particle averaging.

Particle averaging is more widely known through its use in cryo-electron microscopy (cryo-EM), where it has been used with great success to derive protein structures at very high resolution ^2,3,4^. Particles (single observations of a structure in 2D) are classified, aligned (rotation and translation) and those determined to be of sufficient quality are averaged. This data enables a high resolution reconstruction of the structure of the protein, constrained by knowledge of its constitutive components.

A similar approach can be taken in fluorescence microscopy ^5^, with a popular example being the nuclear pore complex (NPC), a large molecular complex inserted in the nuclear envelope in eukaryotic cells that allows exchange between the nucleus and cytoplasm ^6^. NPCs are an attractive target as they are well-characterised structures with the most common target, nup96, having only 32 labelling sites, and they are oriented consistently relative to the nuclear membrane (the centre of which is horizontal for adherent cells). Particle averaging of the nuclear pore complex ^7^, adapted to benefit from 3D data, improves resolution ^8,9,10,11^, and this approach has also been applied to DNA origami ^12^, centrioles ^13^ and ciliary distal appendages ^14^.

Current particle averaging approaches in fluorescence microscopy suffer from two major challenges. First, aligning image patches to each other is algorithmically quadratic in the number of images, and linear in the number of pixels, limiting the method to small, simple structures (a few tens of fluorophores and on a scale of 100nm) ^8,13,9,10^ even when steps are taken to make it more efficient ^15^. This issue can be avoided to some extent by constraining the fit to parametrised models (e.g. line segments) ^16,17,15,14^ but this limits the type of structures being fitted and is still very slow, for example taking 20 hours to fit a 5.2 μm section of microtubule as a tube ^15^. All these approaches to 3D rotation fitting are prone to becoming stuck in local minima, meaning that optimisation may often not be possible. This problem becomes more severe with complex structures. Rather than fusing particles, patches can be compared to attempt to detect self-similar structures ^18^, though this means the analysis does not gain any resolution by combining information from the different patches.

The second major challenge is that at the lengthscales accessible to fluorescence microscopy, almost all structures of interest are heterogeneous. To mitigate this, particles can be separated into different classes which are thought to correspond to different structures ^19^, and limited continuous heterogeneity can be characterised in the nuclear pore complex, when subject to small deformations ^20^. However, neither of these approaches can deal with the large, continuous heterogeneities present in most biological systems.

Here we describe Simultaneous QUAntification of Structure and Structural Heterogeneity (SQUASSH), an approach to simultaneously fit both the structure and heterogeneity in 3D fluorescence microscopy images of the same biological structure. Our approach improves the reliability of alignment and the optimisation speed by using a deep learning system to find the orientation and position in each observation. Such systems can learn the underlying manifold ^21^, and recent smooth descriptors of 3D rotation ^22^ have helped to improve performance, allowing a 3D structure to be extracted from 2D observations of structures ^23^.

Heterogeneity here covers any type of variation which can be easily parametrised, and includes repeating structures within a single field of view, stretch, scaling, and various types of deformation. Quantifying the structure and variability of biological structures at the same time offers the ability to dissect different sources of noise (biological variation, random experimental noise and systematic experimental biases), promising higher accuracy and new biological discoveries.

## 2 Results

SQUASSH analysis uses a convolutional neural network (CNN) to provide the rotation and translation between the current structural model and each particular instance of data. This has two major benefits: the system can recognise repeated observations at similar angles (meaning that the problem is not quadratic with the number of images as for pairwise alignment), and it can align much more complex structures. Our tests of the optimisation of the rotation fitting (see Supplementary Video 1) give confidence that the network is learning the underlying manifold.

In fluorescence microscopy, fluorophores are rendered into an image with an approximately Gaussian point spread function (e.g. confocal), or recorded as a list of fluorophore positions (e.g., single molecule localization microscopy (SMLM)). We developed a fully differentiable rendering pipeline that simulates the imaging process and allows comparison of the expected output of hypothesised fluorophore positions to the data (see Fig. 1). For SMLM data, this pipeline is used to render the localised points that constitute the data, while for other data types the images are used directly. In both cases the rendering pipeline is used to create images from the structural model, which can be compared to the experimental data. The structural model starts with a randomly generated cloud of points in 3D space and then optimises the positions (see Supplementary Video 2).

**Fig. 1.**
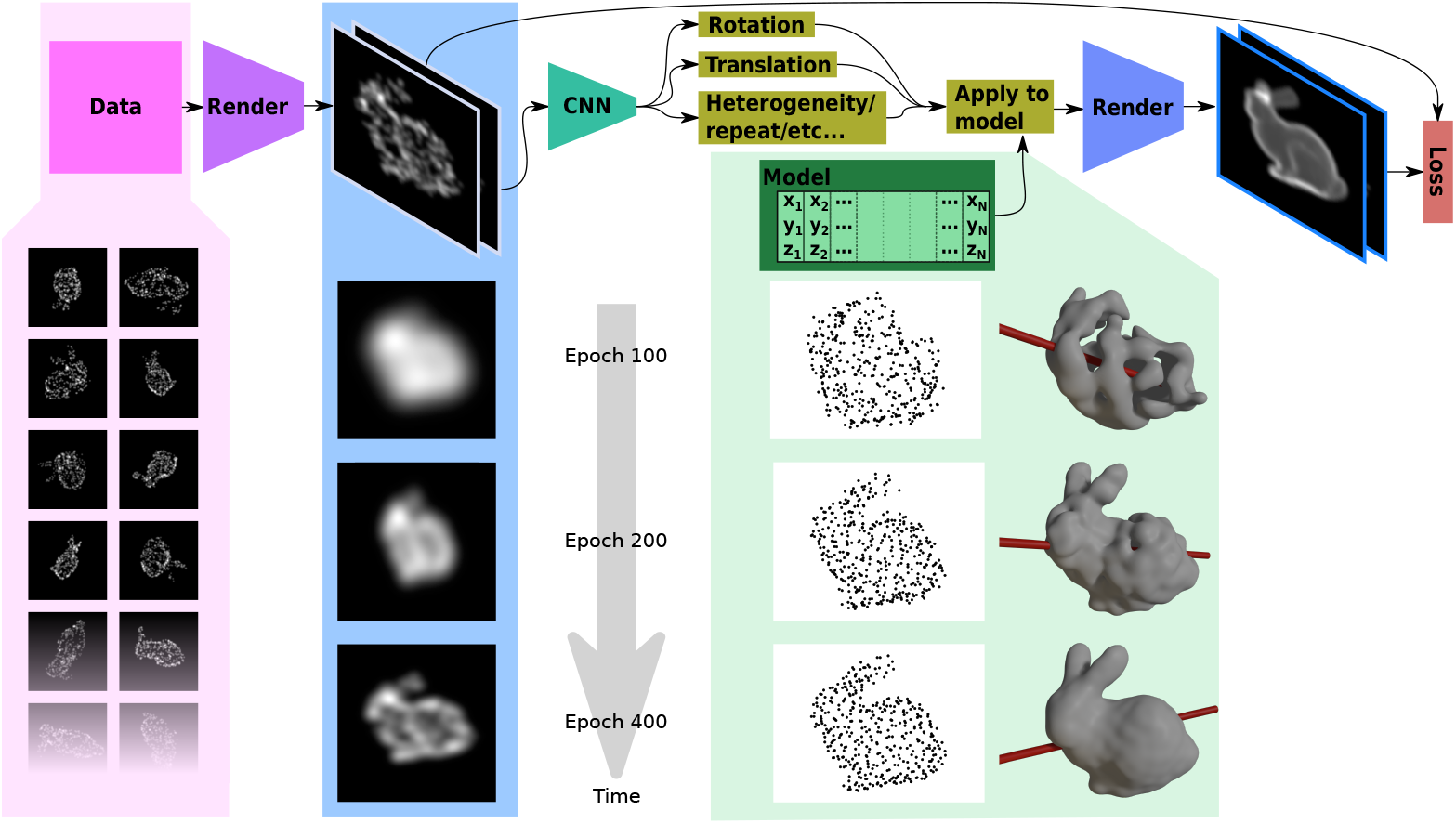
A neural network with heterogeneity parametrisation enables model-free fitting of fluorescence microscopy data. The SQUASSH approach is demonstrated on simulated SMLM data of the Stanford Bunny ^24^, with random stretches along one axis added for heterogeneity. Data (pink highlight area) consists of either 3D localisation microscopy data or fluorescence microscopy image stacks of different instances of a single structure type. For each instance of the structure multiple slice images of the data are rendered (blue highlight area). The rendering blur is decreased over the course of the optimisation, as seen in the three examples in the lower half of the highlight area. These views pass through a network, yielding a rotation and translation, and parameters corresponding to the heterogeneity. Using these the current structural model, consisting of fluorophores at a number of positions (green box), is rendered with the predicted stretch applied in each case. The error between the rendering of the current structural model using computed parameters and the data is used to update the structural model, the heterogeneity model, and the network weights. Over the course of optimisation (see time arrow and epoch numbers) the blur is decreased, allowing the structure to first be optimised at longer lengthscales before attempting to resolve the detail of the structure. This leads to a gradual improvement in the output structural model and heterogeneity model (see green highlight area, where all points are displayed on the left and an isosurface along with the deduced axis of stretch on the right) and a good agreement with the ground truth structure.

Fluorescence images are sparse, with small intensity gradients across much of the image. This makes the network particularly prone to getting stuck during optimisation. We avoid this by initially rendering with a large Gaussian (or, for a non-SMLM image, blurring the input image with a Gaussian). This means that at the beginning of the optimisation process the large scale structure of the sample is optimised. Over the course of the optimisation the blur is lowered, from coarse to fine, so more detail is visible in the rendered structures, and therefore more details can be found in the structural model ^25^ (see Fig. 1, blue panel).

The other major challenge is that from the nanoscale up, almost all biological structures vary from one instance/observation to another. Here we allow the user to build in a parametrised description of the structural heterogeneity of the dataset (see Supplementary Fig. 1). For continuous variation this could be size, squash/stretch, or other types of deformation. Where the biological system being imaged may have many repeating units within a field of view, such as with the repeating units of the microtubule or spectrin rings in axons, the number of repeats in a field of view can be specified as a parameter to be fitted. Allowing both the optimisation of discrete and continuous types of variability provides an important new capability for analysing biological data.

The underlying base structure of the sample and a quantification of how the structure varies from image to image are found as the CNN and structural model are optimised (see Fig. 1, green panel). It should be noted that we are not training the network to use it again: it is the process of training the network which yields the base structure, rotations and heterogeneity, and much like the adversary in GANs (generative adversarial networks) ^26^, the network is discarded after training.

### 2.1 Modelling of structure and heterogeneity in SMLM data reveals experimental effects at the nanoscale

As a first demonstration of the capability of SQUASSH we used two SMLM datasets of nuclear pore complexes (NPCs), a popular exemplar structure for particle averaging ^7,27^ due to the presence of hundreds of repeats of the structure in a single nucleus. The protein Nup96 is distributed across two rings (rotated relative to each other) of diameter 108 nm separated by 57 nm, each of which are decorated with eight pairs of proteins ^28,29^. The first dataset was imaged using resolution-enhanced sequential imaging (RESI), with measurements indicating a lateral precision of 1 nm, with the axial precision not specified ^1^. The second was imaged using 4Pi-STORM ^30^ (Stochastic Optical Reconstruction Microscopy), an interferometric technique expected to have an out-of-plane resolution roughly double the in-plane resolution.

For each dataset SQUASSH derived a structural model for Nup96 from randomly initialised positions (see Supplementary Video 3). Heterogeneity was added to the structural model in the form of an overall scale and a single axis along which the structure can squash or stretch. The angle of the axis relative to the structural model is optimised during training, and the network makes a prediction of the degree of stretch or squash in each instance. This stretch along an axis would allow either a squashing of the ring, as previously hypothesised ^20^ to be the dominant mode of deformation, or movement of the two rings relative to each other (or some combination of both).

The resulting structure agrees closely with EM and previous high resolution particle averaging results. Each ring displays eight motifs, and for RESI showing a clear dumbell shape indicating the two Nup96 sites, similar to results seen from particle averaging techniques ^11,1^. The median structure for RESI had a ring diameter of 113 ± 1 nm and ring spacing of 57.5 ± 1.2 nm while for 4Pi-STORM the median structure had a diameter of 109 ± 2 nm and ring spacing of 63.2 ± 2.9 nm (errors at 1*σ* over 10 runs). For both datasets the stretch axis was found to be along the barrel of the nuclear pore. The derived structure and distribution of stretch across the population is shown for the RESI dataset in Figure 2a and b, while the structure for the 4Pi-STORM dataset is shown in Figure 2c and d (note Figure 2 shows the result of a single run, histograms of the stretch over multiple runs showing the variability are shown in Supplementary Fig 2).

**Fig. 2.**
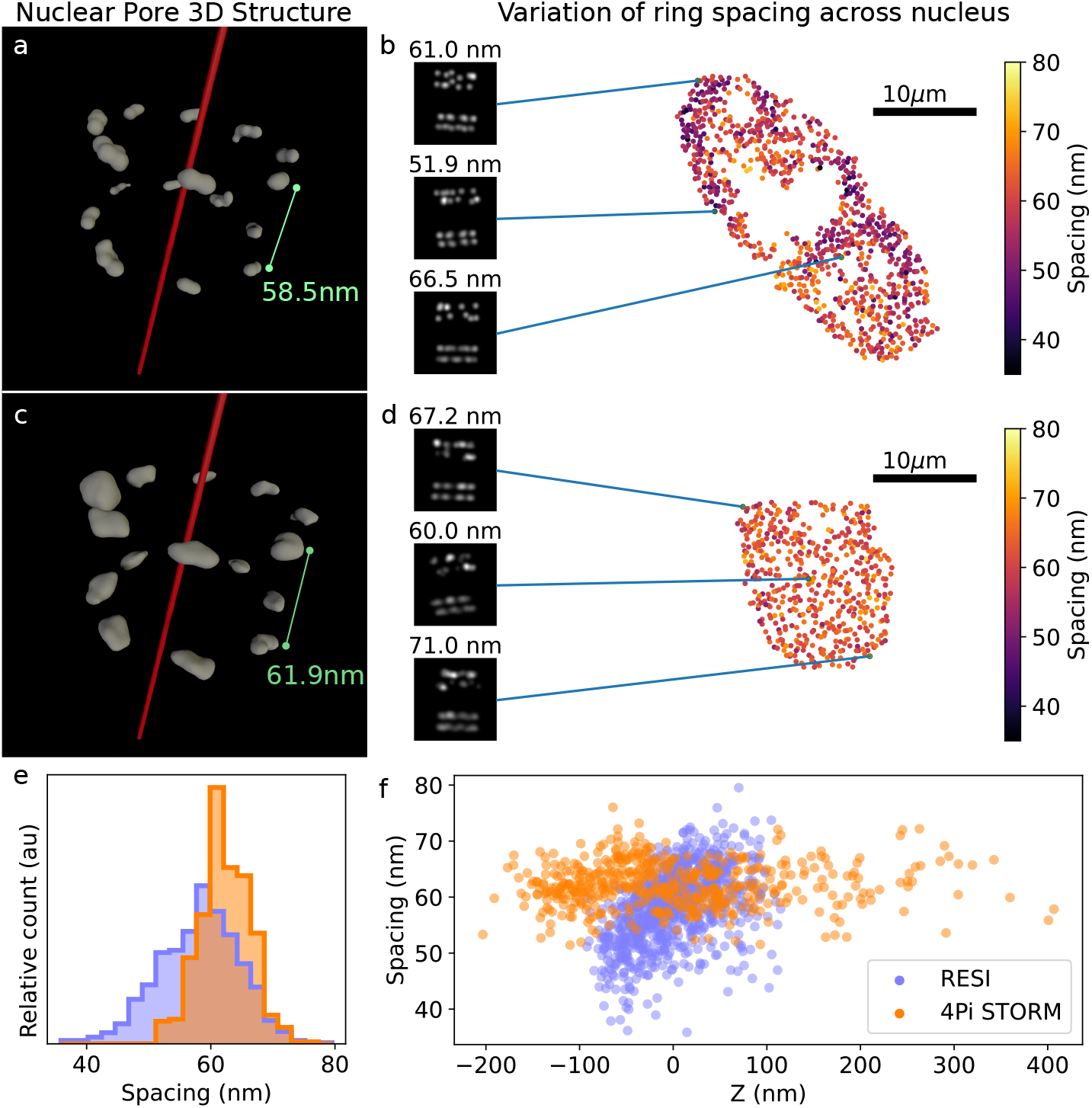
Modelling the structure of nuclear pores with heterogeneity included reveals differences in localisation performance and systematic bias between different nanoscale imaging techniques. (a) Nup96 structure derived from RESI dataset allowing a single axis of stretch which optimised to the axis of the barrel of the nuclear pore complex. The structure displayed is an isosurface. (b) maps the spatial heterogeneity of the stretch of the nuclear pore complex, with side projections of three sampled nuclear pores. Raw data is displayed on top and fitted structural model on the bottom for each pore. (c) Nup96 structure derived from 4Pi-STORM dataset. (d) maps the spatial heterogeneity of a single cell from this dataset. (e) plots the distribution of the ring spacing for the RESI and 4Pi-STORM datasets, demonstrating the larger range of ring spacings observed in the RESI data. (f) spacing of the nuclear pore complex plotted against z position of the complex. The 4Pi-STORM shows a greater range of z positions but no relationship between the z position and the spacing (Pearson’s Correlation Coefficient *ρ*=0.02, p=0.61) while the RESI data shows a clear correlation between z position and ring spacing (*ρ*=0.42, p=10^−55^).

It should be noted that a more precise structure can be obtained by setting the number of fluorophores to the correct value where this is known (see Supplementary Fig. 3). However, for highly symmetric structures the optimisation process will align missing fluorophores which then disappear in the output structural model (see Supplementary Fig. 3), which cannot be avoided without defining the expected symmetry.

As is clear from Figure 2e, the RESI dataset shows a considerably larger degree of stretch than the 4PI-STORM dataset. In the real space mapping of the nuclear pore ring spacing across the nucleus (Figure 2b for RESI and Figure 2d for a sample cell for 4Pi-STORM) the distribution for 4Pi-STORM appears random, but that for RESI is strongly ordered along the shape of the cell. A check of the correlation between the ring spacing factor and the z position (Figure 2f) reveals a strong correlation for the RESI data but none for the 4Pi-STORM (for cell displayed in Fig. 2d). The most plausible explanation for the correlation between z and ring spacing is a systematic bias arising from incomplete modelling of the variation of astigmatism with z ^31^. A similar variation of ring thickness with astigmatism was observed in a previous nuclear pore complex dataset ^27^.

These results demonstrate that SQUASSH can yield results for structures like the nuclear pore complex comparable to current particle averaging techniques, but without the need to impose symmetry constraints. Good results can be achieved even when using only a few hundred images (see Supplementary Fig. 4). Our approach can also identify systematic biases in the data, allowing better assessment of the highest performing microscopy techniques.

### 2.2 SQUASSH analysis enables fluorescence imaging of the 8 nm alpha tubulin repeat in microtubules

Microtubules are cytoskeleton filaments made of alpha/beta tubulin heterodimers. Individual dimers are approximately 8 nm in size and arranged head-to-tail to form protofilaments. Typically 13 protofilaments form a characteristic hollow structure approximately 25 nm in diameter (Figure 3), although in vitro assembled microtubules usually have 14 protofilaments with some variations around this number ^32^.

**Fig. 3.**
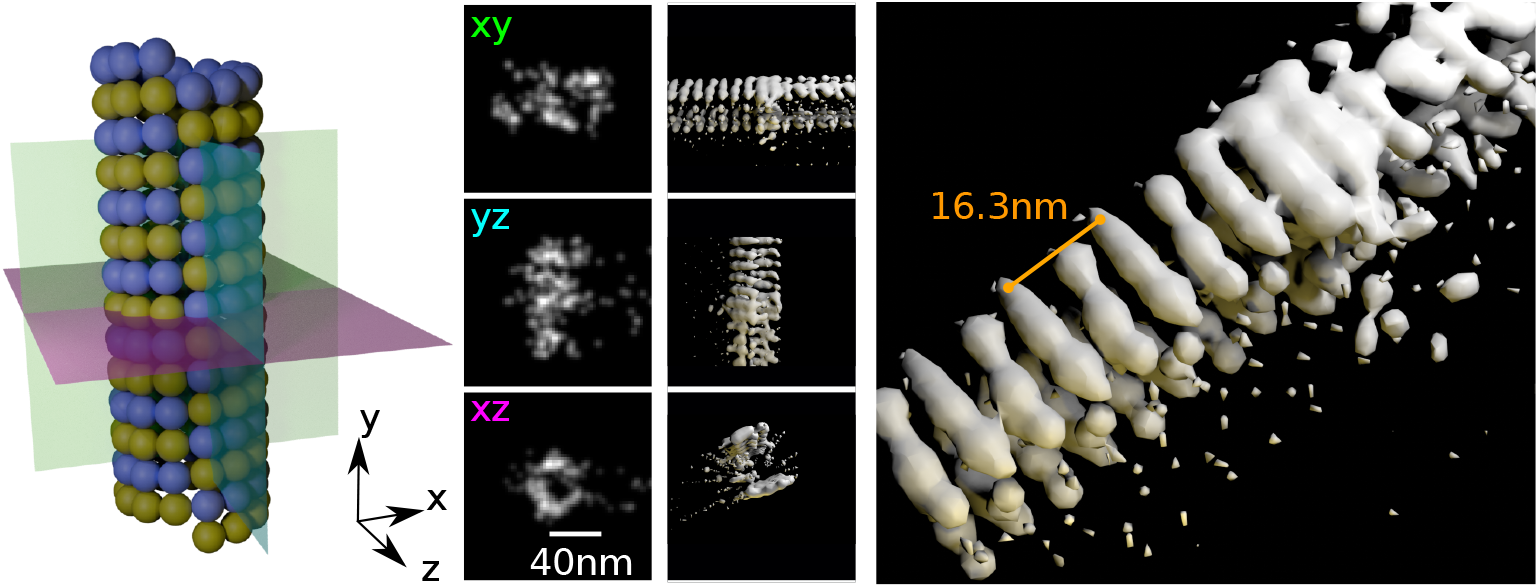
SQUASSH analysis of PAINT data reveals 8 nm alpha-tubulin repeat. *In vitro* microtubules (left panel shows expected structure) were imaged using DNA-PAINT. Microtubules were segmented from the background and were then divided into equal sized patches, with padding to avoid information being close to the edge of the image. Left-middle panel shows example data, with xy, yz and xz projections shown. SQUASSH analysis yields the structure shown from different viewing angles in the right-middle and right panels. The repeat distance of the underlying structural model is 16.3 nm, and the alpha tubulin repeat is clearly visible within the repeating unit as around 8.2 nm.

Using a custom built, low-drift microscope we acquired DNA-PAINT data of in vitro assembled microtubules in which DNA imaging strands were covalently attached to the C-terminal tails of alpha tubulins, and inferred z positions of single molecule localizations from the number of photons emitted using the SIMPLER algorithm ^33^.

Reconstructed 3D images of individual microtubule 3D patches were then broken down into segments and analysed using SQUASSH. The system was parametrised as repeating structure as EM results seem to indicate that spacings along the microtubule are highly regular ^34^ (in the absence of microtubule associated proteins, or other chemicals which change the microtubule structure ^35,36^). An axis of stretch perpendicular to the repetition axis allowed any issues with variation in the z positions found by SIMPLER, similar to those observed for RESI, to be folded into the heterogeneity parametrisation. The results for a repeat segment bounded to be between 14 and 18 nm are shown in Fig. 3. The microtubule structure is found as partial rings (with some information missing at the top and bottom of the microtubule). The system found a repeat distance of 16.3 nm, with two clear repeats separated by around 8.2 nm. This is slightly lower than the average expected separation of 8.4 nm. However, this is not due to an inaccuracy in the fitting, but rather a systematic bias produced by the interaction of the random scatter in fluorophores localised by SMLM and this particular structure. The effect can be seen more severely when fitting with a patch size half as big (64 nm), for which the repeat distance is 7.4 nm (see Supplementary Fig. 5). This effect can be demonstrated by simulating the microtubule structure and measuring the distances present in the data. As the scatter is increased, the observed repeat distance decreases (see Supplementary Fig. 6). It should be noted that the mean repeat distance is not changing: this remains at 8.4 nm, as we confirmed in our experimental data with a Fourier transform of a small number of high quality data patches (see Supplementary Fig. 7). In a repeating structure, scatter causes the *modal* spacing between points to decrease, and in short segments this is better fitted by reducing the spacing of the structural model. This illustrates how efforts to extract local structure must take into account the particular properties of the imaging method.

### 2.3 Extracting fluorophore distribution enables quantification of structure and variation from the nano to the micro scale

To illustrate that SQUASSH heterogeneity models can capture the types of variability that inform on biology we will consider three examples, covering structures with lengthscales from a few hundred nm to just under a mm. These structures are imaged with different fluorescence microscopy techniques including SMLM, lattice light sheet microscopy and confocal microscopy, demonstrating the potential of SQUASSH analysis to apply to all fluorescence imaging modalities, and over the wide lengthscale of fluorescence microscopy.

Firstly we will examine how SQUASSH can be used to uncover correlations in the structural variation of spectrin rings within the axons of neurons, which are known to be organised with a periodicity of around 190 nm ^37,38^. We analysed SMLM data of spectrin rings, using SQUASSH to consider each patch as arising from a number of repeats of an unknown structure. Heterogeneity was introduced with a stretch along the structural model axis and a circular harmonic distortion (intuitively, a squash which may also change the shape) perpendicular to the model axis. Both distortions were constant across a single image patch, which contained around 5-7 spectrin rings. We were able to retrieve a base structure (see Fig. 4), a ring which is subject to various types of distortion. The spacing between the rings has an average value of 194 nm, consistent with previous measurements, and the variation in spacing across the sample can be mapped (see Supplementary Fig. 8). The distortion of the rings can be characterised using PCA, with the first three PCA components shown in Fig. 4 roughly corresponding to scale, keystoning and stretch/squash (see also Supplementary Video 4). By taking the central second moments of the data, parameters corresponding to the width and height of the spectrin ring could be obtained. By comparing the width and aspect ratio of the observed rings across all patches (see Fig. 4), we revealed a strong correlation between increased width and increased aspect ratio. As might be expected intuitively, for axons plated on a coverslip as the axons get bigger they become more squashed.

**Fig. 4.**
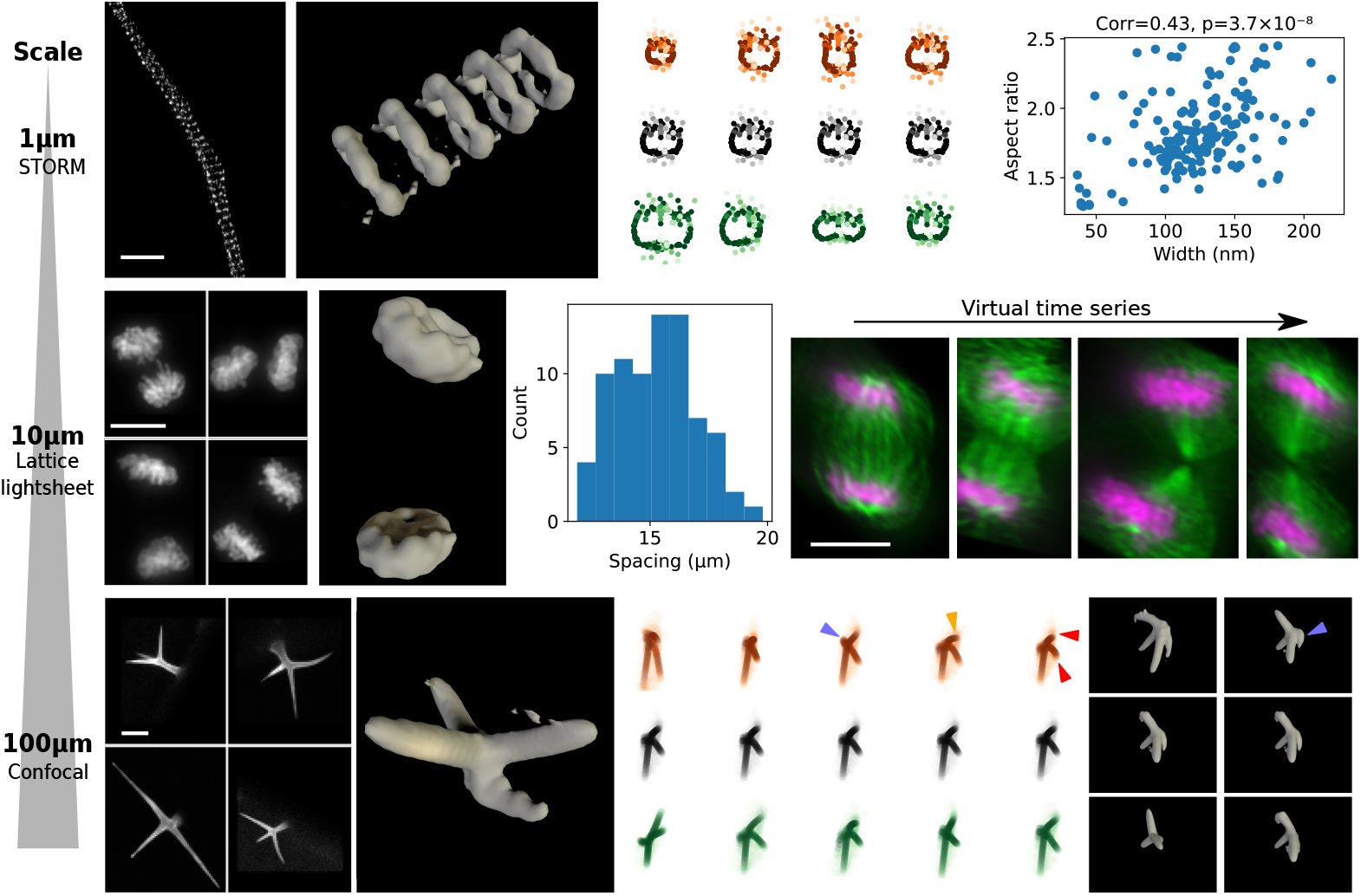
SQUASSH analysis can reveal structure and enable analysis from the nanoscale up to the mm scale. The scalebar length for each row is given in the leftmost coulmn. Row one: spectrin rings in axons, imaged using SMLM (panel 1) can be analysed to reveal the repeating structure of spectrin rings (panel 2) along with the variation of modes of distortion. Panel 3 shows the first four PCA components, with the mean structure in black and the structure one standard deviation from the mean shown in orange and green. Panel 4 shows that by plotting the x axis size (given by the standard deviation of the points in x) against the aspect ratio (the ratio of the x and z size), it can be determined that wider axons are likely to have a higher aspect ratio. Row two: dividing fixed cells in telophase imaged using lattice light sheet microscopy (panel 1) were modelled as two structures with a variable separation and angle. SQUASSH analysis reproduces the two chromatin plates (panel 2). Plotting the separation of the structures per image (panel 3) allows observation of the relative frequency of different separation distances, and re-ordering the dataset according to separation distance allows us to create a virtual dividing cell with chromatin shown in magenta and tubulin in green. Row 3: Plant trichomes imaged using fluorescence microscopy (panel 1) analysed using SQUASSH analysis yield a trichome structural model (panel 2) and allow the identification using PCA of primary modes of variation (panel 3). Bending of the trichome branches and neck is highlighted by coloured arrows. The two columns of panel 4 show 3D renderings of the first and third columns of panel 3.

Secondly, we investigated how quantification of structural heterogeneity can allow reconstruction of a virtual time-varying event from fixed cell data. In a dataset of fixed cell lattice light sheet images of cells in telophase ^39^, the two plates of chromatin could be seen to be separated at variable distances. To analyse the data it was downsampled, as details of the chromatin structure are not easily parametrisable, and modelled as two copies, separated by a variable distance and angle. After fitting it is clear that the reconstruction reproduces the general shape of the chromatin well, with the axis angle and variable separation allowing fitting at different stages of telophase. The separation of the two copies can be used to create a virtual dividing cell from the fixed cell data (see Supplementary Video 5).

Thirdly, we analysed data of trichomes. Trichomes in Arabidopsis thaliana are single-celled epidermal protrusions on the adaxial (upper) side of the leaf, with a neck and three to four tapering branches. The whole structure is around 300 *μ*m in size. The leaves were stained and imaged using fluorescence confocal microscopy. For SQUASSH analysis this was modelled as a structure with a distortion applied. The distortion was created using one vector for each corner of the cube, with the distortion at each point extrapolated between them (see Supplementary Fig. 9). This parametrisation allows distortions such as variability in angles between branches and in relative branch lengths. Fig. 4 shows the structure of the trichomes, which is retrieved despite the highly unconstrained parametrisation of both the structural model and the distortion. The primary components of variation are dominated by size, relative length of the branches, and angle between the branches. In some higher order moments some more subtle effects can be seen, such as a bend in a branch, probably arising from a number of trichomes in which a branch was kinked as a result of sample preparation. Thus our trichome results demonstrate that our approach can work for very generalised heterogeneity parametrisation, and for very large structures.

## 3 Discussion

By modelling the image formation process and the variability of biological structure we have been able to carry out a completely free fitting of structure and a moderately constrained fitting of heterogeneity simultaneously, which we have demonstrated by recreating the standard test structure of the nuclear pore complex, observing the 8 nm repeat of *α*-tubulin in microtubules with fluorescence microscopy for the first time, and by enabling automated quantification and parametrisation of the structure and heterogeneity of structures from the nm to the mm scale. This represents a step change in capability on a number of fronts. Our approach requires only a small alteration in the data pre-processing to accommodate different fluorescence microscopy techniques, making it applicable to a very wide variety of datasets. Since in SQUASSH the pose fitting and structure are learned for each specific dataset, no prior structural information is built in, in contrast to most other deep-learning based approaches. This means that we do not have the common drawback of deep learning based approaches, that what is returned in an image is strongly influenced by the training data in ways difficult for a user to predict or detect. Our model of heterogeneity has been demonstrated to accommodate a number of biologically relevant types of variation in the current study, from repeating structures to multiple types of deformation. The coding system is simple enough that creating an additional heterogeneity model requires only a single python program. In the future, this approach could be used on a wide variety of biological structures across scales, allowing direct extraction of structure, heterogneity and dynamics, and enabling discovery from nanoscale characterisation of macromolecular complexes to the systems biology of development.

## 4 Methods

### 4.1 General software structure

SQUASSH fits a 3D structural model in the form of a point cloud to a set of data. An overview architecture for this is shown in Fig. 5. The quality of fit is measured (and thereby optimized) by rendering the structural model into a series of 2D images that correspond to the data and then comparing these to the data. In order for the comparison to be meaningful, the structural model needs to be rendered at the correct pose, i.e. position (*t*) and rotation (*R*), and these are found using a deep neural network. For each datum (either a point cloud representing a single instance of the structure under investigation, or a 3D image stack of a single structure), all the images (where these images all represent a single structure) are processed by a convolution network (CNN), the features are concatenated, then further processed by a fully connected network (FCN), which outputs the pose. The network and 3D point cloud are then optimized together.

**Fig. 5.**
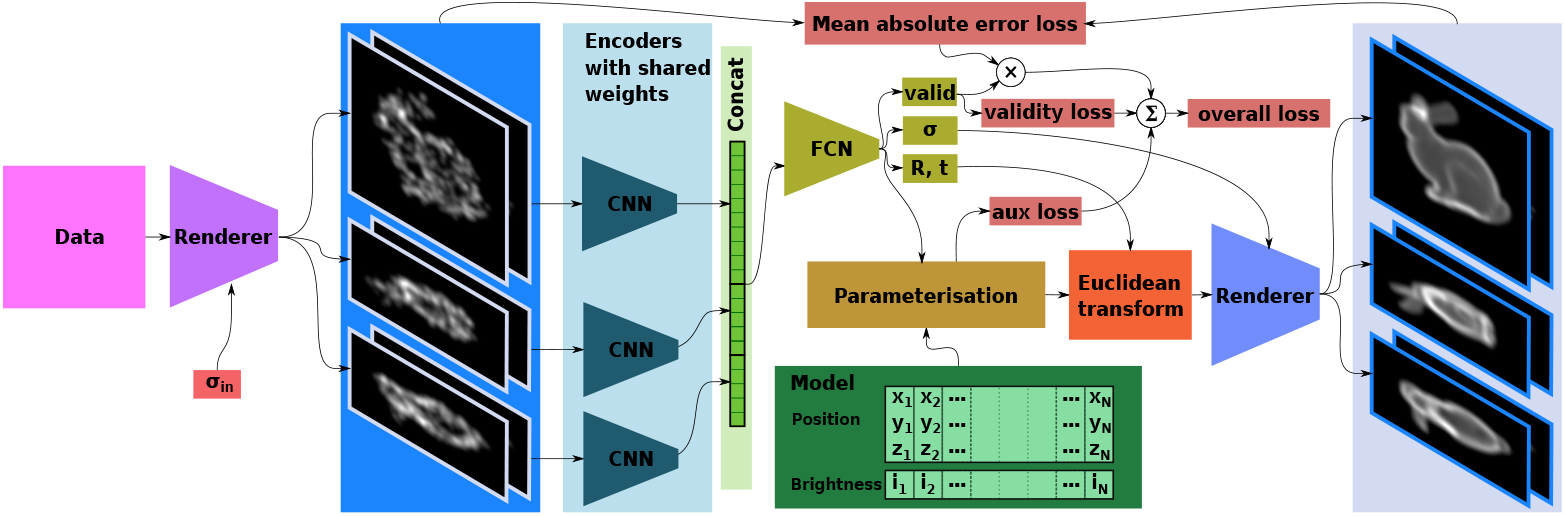
Network structure for SQUASSH showing heterogeneity parametrisation, pose and validity loss. Validity loss enables poor quality data to be automatically excluded and thus not contribute to the structure. Data enters on the left and is rendered with to a specified resolution into a number of images based around projections along the 3 Euclidean axes. Since resolution is usually lower in *Z*, the rendered images need not be isotropic. The images are then processed with a CNN encoder, then an FCN

There are a number of architectural details needed to make a practical system. The first step renders the input data into a series of 2D images (for example XY, XZ, YZ projections) with a given level of blur (*σ*_in_). Pointwise data is directly rendered using a Gaussian PSF determined by *σ*_in_, and volumetric data is blurred to give it the equivalent PSF, and projected or sampled as necessary. In many cases, the *Z* resolution of the data is lower than *XY*, so fewer pixels are needed in *Z*, and *σ*_in_ for the *Z* axis may be correspondingly larger.

The network itself is a purposefully simple structure. The encoder has 5 strided conv-BatchNorm ^40^-SiLU ^41^ activation blocks, using 3 × 3 convolutions to increase the channel count and decrease the size by a factor of 2. From input to output, the channel counts goes 1–16– 32–64–128–256. The encoder is applied to all input images and the outputs are flattened and concatenated, with the resulting number of channels, *N*, depending on the number and size of the images.

The concatenated encoder outputs feed into an FCN with a similar structure (FC-BatchNorm-SiLU), with the channel counts going *N*–256–256–*M*, where *M* is the number of output logits. There are *M* ≥ 11 logits, since 3 channels are needed to encode translation, 6 are used to encode a rotation ^42^, one to encode the validity and one used to encode *σ. σ* determines the PSF of the rendering following our previous work ^23^ and allows the system to account for inaccuracies in the data such as localisation errors and scatter of the fluorophore positions. The validity output is a classification of whether a datum is valid, and allows the network to essentially ignore a datum at the penalty of a fixed cost. This adds robustness to the fitting process and reduces the need to very carefully hand select input data.

Finally, the network outputs are limited in range to reasonable values. For example, *σ* cannot drop below *σ*_in_, so it is computed from the logit, *l*_*σ*_ using SoftPlus ^43^ with the formula *σ* = *σ*_in_ (1 softplus (*l*_*σ*_)). If the data is not already centred, we centre the data as much as practical, so we limit translation to ±10% of the patch size giving *t*_*x*_ = 0.1*s* tanh *l*_*x*_, where *s* is the size of the visible patch and *l*_*x*_ is the logit corresponding to the translation in X (*t*_*x*_).

The 3D structural model itself is represented as a collection of 3D points each with an associated brightness between 0 and 1. The brightness allows the optimization to remove points which are unnecessary, and can also allow the structural model to represent a low precision structure with a large number of low brightness points. The 3D positions are stored in scaled length units so that the range of positions is roughly ±0.05, comparable convolution kernel values often found in practice, to avoid the need for different learning rates. Likewise brightness is computed from a stored value, *b* as sigmoid (100*b*).

Such a 3D structural model is inherently limited to being able to represent only single instances or rigid structures, so the network has a parameterisation block, which is specific to the type of structure being modelled. The parameterisation modifies the structural model for each instance of the data according to logits generated by the network. The parameterisation may also have some learnable parameters.

For example, for the nuclear pore complex, the parameterisation consists of a learned axis, and two scaling parameters generated by the network. The 3D structural model is scaled along the axis and isotropically in the plane normal to the axis (these combine to give overall scale). After the parameterisation is applied, the points are transformed using *R* and *t*, then rendered using a Gaussian PSF, with the computed *σ*. Changing this parameterisation allows SQUASSH to deal with very varied data as detailed below.

The loss between each pair of images is computed by normalizing the images so that they sum to 1, then computing the mean absolute error (MAE). The overall loss is the weighted sum of the individual losses. For a parameterisation with scaling factors (such as for nuclear pore data), the auxiliary loss is the absolute value of the logarithm of the scale which helps the average scale to settle to around 1, an effect which is not especially sensitive to the weighting, but aids convergence.

In general, incoming data is not entirely clean, meaning that there are likely to be some items of data which are bad, for example due to selection of data from other structures, segmentation errors (e.g. leading to multiple structures in one image), labelling errors (e.g. causing missing structure) and so on. Since these cannot be modelled the same way as the valid data, SQUASSH may attempt to partially build the invalid items into its structural model which can reduce the overall quality. In order to prevent this, and to avoid the need for the user to excessively clean the data (which could lead to bias), the system also attempts to classify data as valid or invalid.

The classification is done by the network and so is updated during training. If a piece of data is classified as invalid, then it incurs no loss on the appearance of the rendered image, so the optimization will not attempt to include that item of data into the structural model. However, a penalty for excluding an item of data is needed since otherwise the system will exclude all data and incur no loss for the entire dataset. The optimizer therefore has a choice as to whether to include any given datum, and it will choose to do so if any given datum can be well modelled. If it cannot be well modelled, then the most optimal choice is to exclude the datum and incur a fixed penalty.

This is implemented by the network emitting a validity classification which is a number between 0 and 1, given by *v*. The MAE loss is *m* and the fixed loss penalty for invalid data is *w*_*v*_. We use the following equation for the loss:

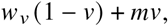

so if *v* = 0 (invalid), we incur a fixed loss and if *v* = 1 (valid) we just use the MAE loss, and smoothly interpolate otherwise.

If *w*_*v*_ is not broadly comparable to the average MAE, then the system will either accept or reject all of the data. In practice the useful range spans roughly a factor of 2. The optimal weighting therefore changes as the optimization progresses and the losses change (especially with changes in *σ*_in_). We compute the weighting dynamically by keeping an exponential moving average of the MAE, and using that to scale the validity loss. Practical values of the weighting are roughly in the range 0.7–1.2, with 0.7 rejecting most of the data and 1.2 accepting most. A value of 2.0 effectively switches off validity checking and uses all of the data. The efficacy of the validity is illustrated in Supplementary Figure 10, where SQUASSH automatically selects nuclear pore complexes with various errors.

We train the network using the Adam ^44^ optimizer over a number of epochs, with both *σ*_in_ and the learning rate starting at a high value and dropping exponentially towards their final values. This allows the network to rapidly acquire the coarse structure, then gradually refine finer and finer structures as the optimization progresses. In practice the change in *σ*_in_ happens once every few epochs to reduce the computational cost of repeatedly rendering the dataset.

The final output is a 3D point cloud, and the parameterisation values for each element of the input data. Recall that the data first needs to be rendered to a series of 2D images. A single 2D projection results in a reflective ambiguity (see Supplemental section 1). While almost any additional information is enough to disambiguate mathematically, there is a tradeoff where more images give more information but are slower to run and more parameters need to be learned in the FCN. A reasonable option is to use 3 projections planes (XY, YZ and XZ). In many cases, a richer representation is useful and we have found that a good balance is struck by a non physical 6 plane rendering, where each of the three projections (XY, YZ, XZ) is split along the projection axis (Z, X, Y respectively), as illustrated in Figure 6. The impact of different rendering methods, localisation precision and labelling level are shown in Supplementary Figures 11 and 12.

**Fig. 6.**
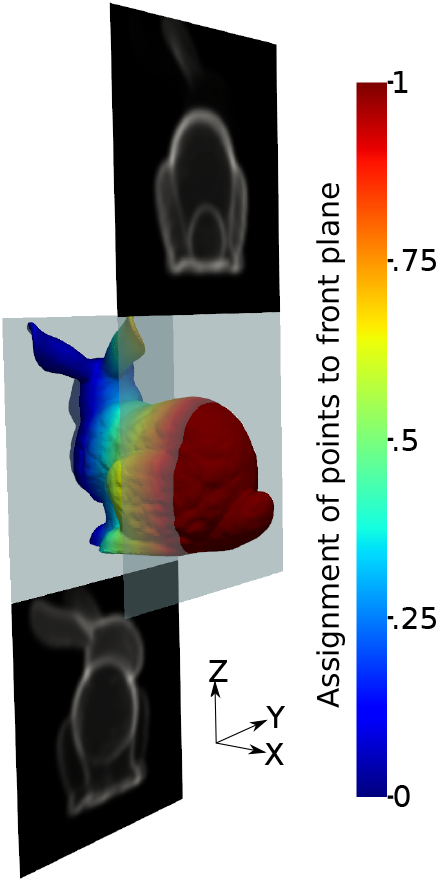
6-plane rendering splits each plane into two along the projection axis. For the YZ projections (shown), there are two YZ planes at different X positions chosen such that 90% of the data lies between the two planes. Each point is rendered in both planes with a brightness weighting according to its X position (illustrated by the colour). The assignment is linear between the planes: if a point is midway between the two planes, the weighting is 0.5 in each plane, if the Z position aligns with one of the planes, its weighting is 1 in that plane and 0 the other. Any points not between the two planes are assigned entirely to the closest plane.

Since the system does learn the underlying manifold, augmentations can help with convergence. However care must be taken to avoid introducing artifacts. In practice we only augment point data, and we do this by performing random rotations about the Z axis.

There is often coupling between various parameters computed by the network, and it assists convergence to remove this where it is practical to do so. For example if the structural model is significantly off centre, then change in rotation will also cause significant motion, so the network needs to learn how R and t couple together. Likewise with an uncentered structural model, scaling will also cause a translation. We have found that in many cases, it improves convergence to centre the structural model. We achieve this by computing the centroid, i.e. the mean of the positions weighted by their intensities, and subtract that off the positions.

All software timings are given for execution on an AMD Ryzen 9 3900X CPU, with an NVidia GeForce RTX 2080Ti GPU.

### 4.2 Nuclear pore complex

#### 4.2.1 Software

The software structure and parameterisation is as described in Section 4.1. To render the data, we use 6 plane rendering with 64 pixels in *X* and *Y*, at 3.9 nm/pixel and 32 in *Z*, at 7.8 nm/pixel, and we use a validity weighting of 1.

To make the training time particularly short, we use a multi stage training process. We start with a small structural model of 35 points, with the rendering FWHM starting at 65nm and reducing to 34nm over 30 epochs, and then down to 13nm over a further 300 epochs, all at a fixed learning rate of 0.0001. We use 8 fold augmentations and since the model is so small, we can use a large batch size (160) for speed.

At this point, we then replace each of the 35 points with 20 new points randomly scattered around its location (giving a total of 700 points). Training then proceeds for 1000 epochs on the data with no augmentations at a FWHM of 13nm, with the learning rate decreasing from 0.0002 to 0.00005 and a batch size of 10.

Training time was approximately 4.8 hours for the RESI data, and 7.7 hours for the 4Pi-STORM data.

#### 4.2.2 Sample preparation and microscopy

The sample preparation and imaging for the RESI data were preformed as described in ^1^. The resulting dataset consists of 1218 nuclear nuclear pores, with an average of 470.2 localisations each from the PAINT imaging, reduced to an average of 12.6 with RESI processing.

Sample preparation and imaging for the 4Pi-STORM data were performed as described in ^30^. The resulting dataset consists of 5822 nuclear pores with an average of 37.2 localisations each.

### 4.3 Microtubules

#### 4.3.1 Software

Unlike nuclear pores, labelling microtubules to separate out individual repeating units would be a very time consuming process, and have to rely on a large amount of human judgement to correctly separate them, which could strongly bias the data. Instead, long sections of microtubules are labelled and then automatically chopped into roughly fixed length segments of around 128 nm which contain a variable number of repeating units. To account for this we use a parameterisation that allows for a variable number of discrete duplicates of the structural model in a fully differentiable manner.

The parameterization needs to be able to generate between *M* and *N* duplicates of the underlying structure. This is done by first duplicating the underlying structure *N* times along a learned axis with a learned spacing. Recall that the structural model contains a list of 3D positions and brightnesses. The parameterisation scales the brigntnesses in each duplicate with a weight between 0 and 1, so a duplicate scaled by 0 is effectively not present. Since the scaling is continuous, the result is differentiable. However independent scaling for each duplicate allows nonsensical structures.

Consider a parameterisation which has between 3 and 6 duplicates. Examples of brightness scalings (*b*_1_ to *b*_6_) for the 6 duplicates would be:

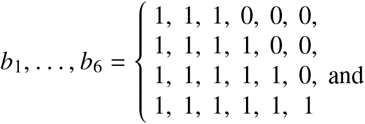

for essentially 3, 4, 5 and 6 repeats. However the sequence 1, 1, 1, 0, 0, 1 would not be valid since that would imply there are three repetitions followed by a large gap followed by a final repetition, which is not a physically realistic structure. In order to ensure that the structure is contiguous, we would like to enforce the constraint that *b*_*i*−*j*_ ≥ *b*_*i*_ for *j* > 0. This means that if *b*_*j*_ = 1, making duplicate *j* be present, then all prior duplicates must be present as well.

In this example, the network must generate 3 outputs, *c*_4_, *c*_5_, *c*_6_ used to compute *b*_4_, *b*_5_, *b*_6_, since *b*_1_, *b*_2_ and *b*_3_ are all 1. Ideally our constraint would be implemented using the max operator, since if *b*_5_ = max (*c*_5_, *c*_6_) and *b*_6_ = *c*_6_, that ensures that *b*_5_ ≥ *b*_6_. However, we cannot use max since it is not differentiable. Instead we use boltz_*β*_, the Boltzmann Operator as defined in ^45^ which behaves as a differentible approximation to the max operator. Our implementation for *M* = 3 and *N* = 6 uses the equations:

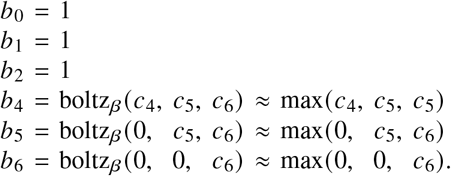

We found *β* = 10 provides a good tradeoff between accuracy of the approximation to max and smoothness for ease of optimization.

The variable duplication can produce structural models which are very off-centre so we perform centering after the parameterisation. The spacing between duplicates is a learned parameter, so like the axis it is the same for all data. Based on our expectations of the underlying structure, we allow the spacing between duplicates to take any value between 14nm and 18nm, and so since the data is chopped into 64 nm chunks we allow for between 3 and 5 repetitions.

Finally, to allow for imperfect calibration of SIMPLER, we allow for a squashing of the structure with can be different for each datum. This requires an additional learnable axis (represented by 3D vector) which is constrained to be orthogonal to the repetition axis using Gram-Schmid orthogonalization. The structural model is then scaled along this axis by at most 20% with the scaling factor *s* given by 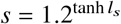, where *l*_*s*_ is the corresponding logit from the network.

Data is rendered using the 6-plane technique, as 64 × 64 for XY and 64 × 32 for XZ and YZ, with an isotropic resolution of 2nm/pixel. The parameterisation allows for between 3 and 5 repetitions of the underlying structural model. We train for 2000 epochs at a batch size of 20 at a learning rate of 0.0001 with the initial and final rendering FWHM going from 20nm to 8nm. The data has 2 fold augmentations and the structural model has 280 points.

The validity weight was 0.6 and training took approximately 11 hours.

#### 4.3.2 Imaging experiments

Porcine brain tubulin was purified by rounds of polymerization/depolymerization as described before ^46^. Digoxigenin-tubulin was obtained by labelling purified tubulin with digoxigenin NHS-ester (Sigma) and purified by cycling following the published protocol ^47^.

Amine-modified DNA oligonucleotides (Integrated DNA Technologies) were attached covalently to C-termini of tubulins using TTL-ligase ^48^. The plasmid used for expression of TTL was a kind gift from Jonathon Howard.

Microtubules were assembled in the presence of 1 mM GMPCPP (Jena Bioscience) immobilized on dichlorodimethylsilane silanized coverslips using Digoxigenin. Flow cells were incubated with a mixture of anti-Digoxigenin-AP, Fab fragments for the attachment of microtubules and Neutravidin for the attachment of biotinylated gold-nanourchins (CytoDiagnostics, GUB10K-100-25) as fiducial markers used to correct for drift in *x*− and *y*−. Coverslips were additionally passivated with beta-casein (Sigma).

Microtubule seeds were prepared by mixing 5 μL 10 mg mL^−1^ DNA-PAINT tubulin, 1 μL 10 mg mL^−1^ digoxygenin-tubulin, 40 μL BRB80, 5 μL 10 mM GMPCPP on ice, incubating for 5 min before placing at 37 °C and incubating for 30 min. A passivated flowcell was incubated with gold biotinylated nano-urchins until approximately 10 - 20 fiducials were observed in a FOV. The flowcell was then washed twice and GMPCPP microtubules were introduced and allowed to bind to Digoxigenin. 0.25% glutaraldehyde in BRB80 was flowed into the chamber and incubated for 5 min before washing with 3 × 100 μL of BRB80 and then 3 × 100 μL of imaging buffer (PBS,1 mM EDTA, 500 mM NaCl, 50 mM glucose).

Imaging was performed on a custom TIRF microscope syste, with 561 nm and 640 nm laser illumination. A TIRF objective (Nikon, CFI Apochromat TIRF 60XC Oil) gave a circular illumination field with diameter 180 μm. Emitted fluorescence was split using a dichroic mirror into two imaging arms using a 4f-configuration, forming the image onto the sensor using achromat lenses (Thorlabs, AC254-200-A) and combining the imaging beams with a knife-edge mirror prism (Thorlabs, MRAK25-P01). To maintain focus throughout long acquisitions a focus-lock module was used. A back-reflected beam from an infrared laser diode was reflected using a 50:50 beamsplitter (Thorlabs, BS013) and imaged onto a camera sensor (Thorlabs, DCC1545M). Movement in the back-reflected spot was detected using custom Python scripts and used to correct for drift in the sample focus by updating the position of the piezo stage on the sample stage (Applied Scientific Instrumentation, PZ-2000 XYZ).

Imaging was performed using 100 pM of Cy3-labelled imaging strands in imaging buffer with Trolox and Pyranose Oxidase/Catalase using maximum laser power (approximately 0.5 kW cm^−2^), acquiring 150,000 frames with an exposure time of 100 ms. Extraction of molecule positions was performed in Picasso and using custom Python scripts.

### 4.4 Spectrin rings in neurons

#### 4.4.1 Software

The parameterisation is an generalisation of the parameterisation described in section 4.3.1. Firstly, since the spacing of the rings varies, we alllow the spacing to vary per-image from between 160 and 240 nm. Second, we have scaling (from 0.5 to 2) which applies orthogonal to the axis of repetition. Finally instead of a simple squash, we apply a more general distortion function orthogonal to the axis of repetition. The data is rendered using 6 plane rendering.

The distortion function is applied radially, that is the vector from a given point to the axis of repetition is computed, and then the point is moved by scaling the the vector. The size of the scaling varies with angle around the axis. The angular variation is parameterized by a number (4) of scalings (of up to 50%) computed equally spaced angles, which are then interpolated to find scalings at any angle. Since the scalings are over a circular domain, we take the discrete Fourier transform of the scalings to find a Fourier series, which can then be evaluated for arbitrary angles. Note that this radial distortion function is applied after centering the structural model, and we centre the model again after the distortion is applied.

The data is rendered into images with 64 pixels in X and Y and 32 in Z, with a resolution of 25 nm/pixel. Training is for 2000 epochs with a batch size of 40, at a learning rate of 0.0001 with the rendering FWHM dropping from 250 to 45nm. We train with 15 augmentations and a structural model size of 300 points. The validity weight is 0.8.

Training took approximately 2.7 hours.

#### 4.4.2 Sample preparation and microscopy

Samples were prepared and data collected as described in ^49^. Well separated neurons were marked up manually, then automatically cut into segments of approximately 1 micron in length. This yields 154 segments with an average 10,700 localisations each from roughly 800 spectrin rings.

### 4.5 Telophase

#### 4.5.1 Software

The is in the form of image stacks. These are cropped around the centroid to produce stacks of 128 × 128 × 64, which are then scaled using average pooling down to 32 × 32 × 32, yielding a final sampling of 864nm/pix in plane and 800nm/pix on the Z axis. For training, the data stacks are rendered into three projections (XY, XZ, YZ), by summing along the Z, Y and X axes.

The parameterisation has a learned axis, which dictates how the nuclei are separated. The structural model is first scaled normal to the axis by up to 50% (i.e. between 0.666 and 1.5), then duplicated and the duplicate is translated along the axis by between 8 and 24*μ*m. The translated duplicate is then rotated around its centre around the two axes orthodonal to the direction of translation, by up to ±45°.

Training proceeds in two stages. The first stage runs for 1000 epochs at a learning rate of 0.0001 with the FWHM dropping from 10*μ*m to 5*μ*m, with no scaling and rotation. During this stage, sice there are only 79 data cubes, we use a tiny batch size (4) to maximize the benefit of stochastic gadient descent’s ability to escape local minima. The second stage introduces rotation and scaling and runs for 10,000 epochs with the FWHM dropping to 500nm and the learning rate decaying to 0.00001.

We set the validity weight to a high value (1000), which prevents the system from rejecting any of the data.

Training took approximately 19 minutes.

#### 4.5.2 Data source

Data was associated with ^39^. The data consists of 79 stacks of 150 z slices, with x and y sizes varing from between 133 and 311 pixels. The sampling is 108 nm/pix in plane and 200 nm/pix in z.

### 4.6 Trichomes

#### 4.6.1 Software

Image stacks of individual trichomes were captured (see Section 4.6.3). Since the trichome neck connects to the leaf, there is often a considerable amount of leaf visible which we do not attempt to model. We have a manual markup step where a 3D plane is provided and all data in the data cube on one side of the plane is set to zero. This is used to cut off the leaf from the neck. Due to the high amount of shape variablility, we do not perform centering since that can cause some features to be moved outside of the visible area. As a result we allow the network to predict larger translations, up to ±30% of the size of the image. Images are downsampled by a factor of 2 in X and Y using average pooling and then images are padded with zeros where necessary to a size of 89 × 89 × 60.

The uncentred data causes an additional difficulty. Since we use a 6 plane rendering, and the data is uncentered, occasionally a rendering plane is essentially empty and contains no data. The per-image normalization process then amplifies the nonexistent data, essentially noise for the data and numerical round-off errors for the rendered structural model. Instead we perform normalization across the entire group of images so the sum of all pixels across all planes sum to 1.

The distortions of trichomes are large and we do not have any particular expectations of what it should be, so we use a general-purpose model of distortion, which is illustrated in Fig. 9. The distortion model operates on a cube, for which we use the smallest cube encompassing the cuboidal volume of data that we are processing with SQUASSH.

The parameterisation is in the form of a vector field defined within that cube. For each point in the structural model there is therefore a corresponding vector from the vector field, and we distort the structural model by adding that vector to the point’s position. This field is parameterised using one 3D vector at each of the eight corners of the cube, giving 24 parameters. Vectors at arbitrary positions within the cube are found using trilinear interpolation of the X, Y and Z components of the vectors.

For a practical implementation, a number of steps are required. Firstly, the X, Y, and Z components are sigmoided network logits, limited independently to 1.2× the size of the cube. Secondly, it is necessary to centre the structural model before the distortion is applied. Consider a case where the structural model is limited to the bottom left 2 × 2 squares in corner of the grid in Fig. 9. The distortion of those is dominated by the bottom left, top left and bottom right vectors, with very little contribution from the top right vector. Finally, the size of the trichomes varies quite strongly, so to prevent the distortion model having to accout for that in addition to actual distortions, we apply an overall scale of up to a factor of 50% (i.e. from 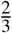 to 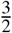) to the structural model after the distortion.

Due the the high degree of variability in the structure, we use a two stage training process. We train first for 16,000 epochs with a learning rate of 0.0001, and a batch size of 16, with the FWHM dropping from 500 to 10*μ*m. During this phase, the complex distortion is disabled and only scaling is permitted. The second stage enables the distortion and proceeds for a further 8,000 epochs with an FWHM of 10*μ*m.

Training took approximately 45 minutes.

#### 4.6.2 Arabidopsis growth and maintenance

*Arabidopsis thaliana (L*.*) Heynh, Columbia-0* seeds were sown on Westland Multi-Purpose John Innes compost. and maintained in a controlled growth chamber at 25°C with a 12 h light /12 hour dark photoperiod. Trichome images were captured from fully expanded leaves of 4-week-old Arabidopsis plants.

#### 4.6.3 Propidium Iodide staining and laser scanning confocal microscopy

Arabidopsis leaves were excised using a razor blade, immersed in a 50 μg/mL working solution of propidium iodide for 10 minutes before being placed, adaxial side up, in the well of a polydimethylsiloxane gasket made on a microscope slide. The well was filled with perfluoroperhydrophenanthrene according to the method of Littlejohn *et al* ^50^. The sample was covered with a coverslip prior to imaging, taking care not to crush trichomes. Trichome images were captured using a Zeiss LSM 880 microscope equipped with Plan-Apochromat 20X/0.8 M27 objective lens, and a 514 nm laser which was used for sample excitation. Emission was collected in the wavelength range of 566–664 nm with gain set to eliminate saturated pixels across each z-stack. 114 Z-stacks were obtained with a pixel size of 3.02 μm in-plane and 5 μm out-of-plane.

## Acknowledgments

We would like to thank Ralf Jungmann, Susanne Reinhardt, Wesley Legant and Yu Shi for providing and assisting with their data. This project was supported by the BBSRC, grant BB/T011823/1, (SuC) and the Royal Society University Research Fellowship scheme, URF\R1\211329 (SiC).

## Author contributions

Approach was designed by ER, SuC and DS, and implemented by ER. Microtubule experiments were performed by DS and L-YC, trichome experiments were perfomed by KW, GL, KB, GMcC, spectrin experiments were perfomed by CL and nuclear pore experiments were performed by MB. Evaluation was performed by SuC, ER, SiC, QC and MM, with input from all other authors on the results of their experiments. ER and SuC wrote the paper and all authors reviewed the paper.

## Data and materials availability

Software and data associated with the paper are available from: https://github.com/edrosten/SQUASSH

## Supplementary Materials

### Supplementary Note

#### 1 Reflection/rotation ambiguity

We reconstruct 3D shapes by predicting Euclidean transformations for a 3D structural model. If the reconstruction uses only one projection for each predicted pose, then the shape can only be determined up to a reflection. Proof:

Consider an abitrary point **X** = [*αX, αY, αZ, α*], **X** ∈ ℙ^3^. For simplicity but without loss of generality we will consider only *α* = 1. The point is transformed by and a Euclidean transform **E**: ℙ^3^ → ℙ^3^:

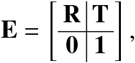

where **R** and **T** are a rotation and translation in R^3^. With affine projection (how microscopes work), the 2D position, **u**, is:

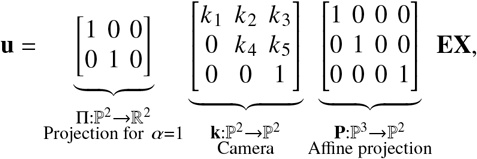

where **k** is an arbitrary camera calibration matrix. Composing the projections and camera yielding **K** = Π**kP** gives:

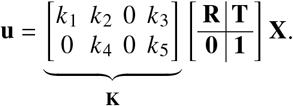

Defining **X**′ = [−*X*, −*Y*, −*Z*, 1] as the reflection of X. then it is clear that:

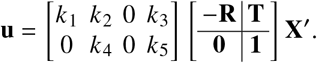

That is not a valid Eucildean transform, bit since **K** has a zero third column, the result is invariant to changes in the third row of the rotation matrix, therefore we can pick 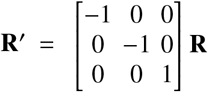, which also gives:

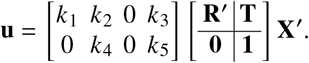

Therefore if we mirror the structure and rotate the pose by 180° around the Z azis, the projected image will be identical.

It is important to note that this holds only because the third columns of **k** is zero, allowing us to substitute a rotation matrix which mirrors two of the axes. This does not hold if there are projections along multiple different axes.

**Fig. 1.**
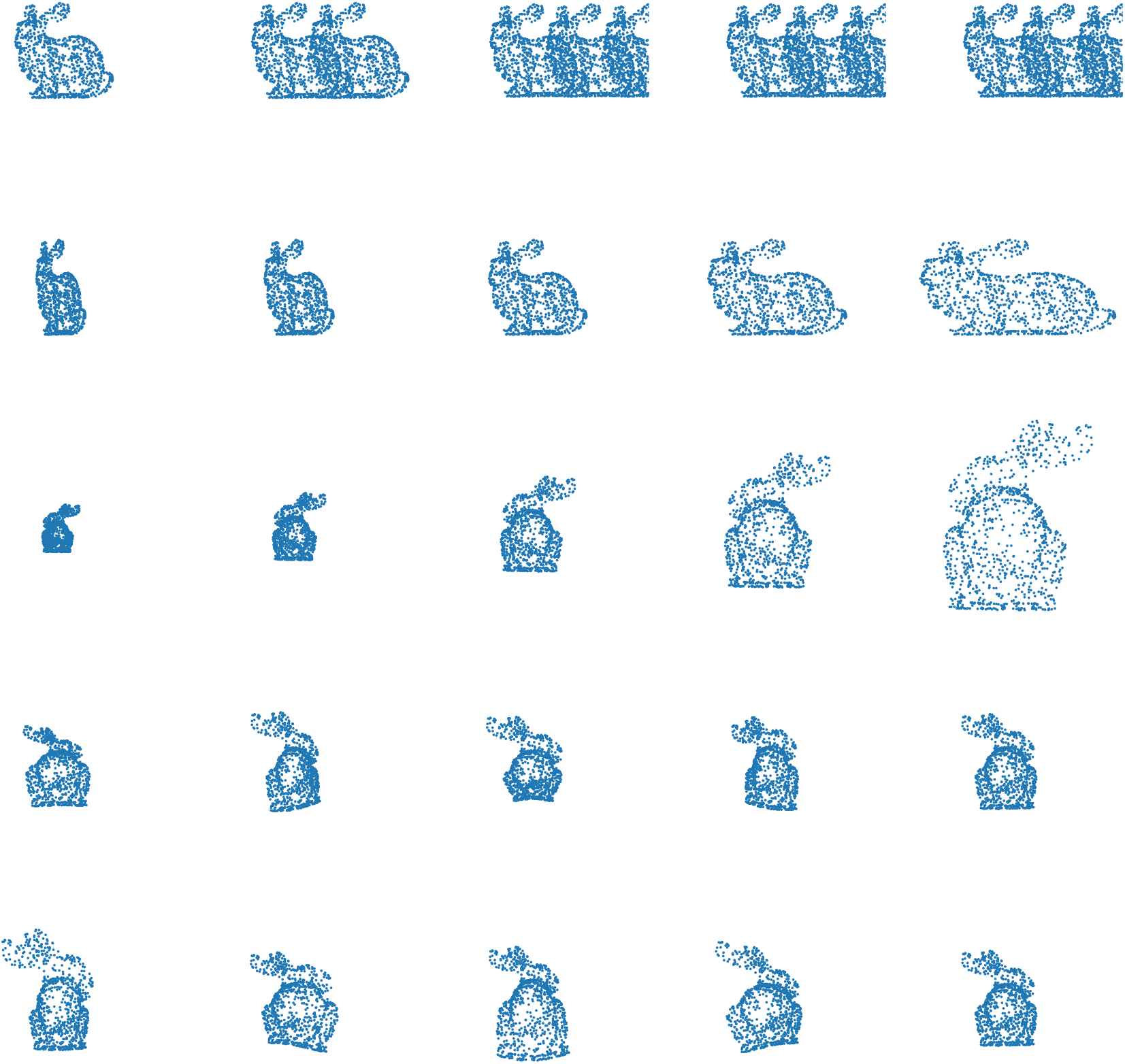
Illustration of different heterogeneity parameterisations. Following Fig. 1 we use the Stanford Bunny as an exemplar structural model, with the primary axis aligned with x. Row 1: variable duplication along an axis. Row 2: elongation along an exis. Row 3: expansion orthongonal to an axis. Rows 4 and 5: various different angle dependent scaling orthogonal to an axis.

**Fig. 2.**
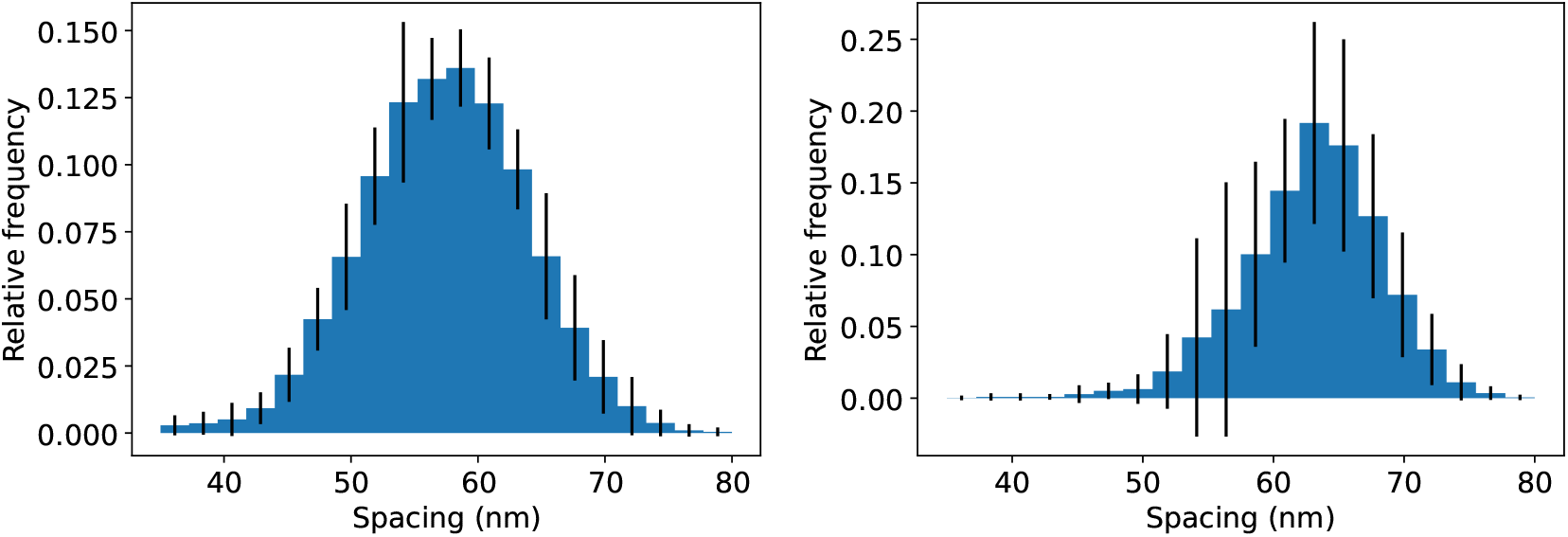
Mean histograms of nuclear pore complesx ring spacings from the RESI dataset (left) and the 4Pi-STORM dataset (right). SQUASSH was rerun 10 times, in order to compute the mean value of histogram bins along with error bars as ±1 standard deviation.

**Fig. 3.**
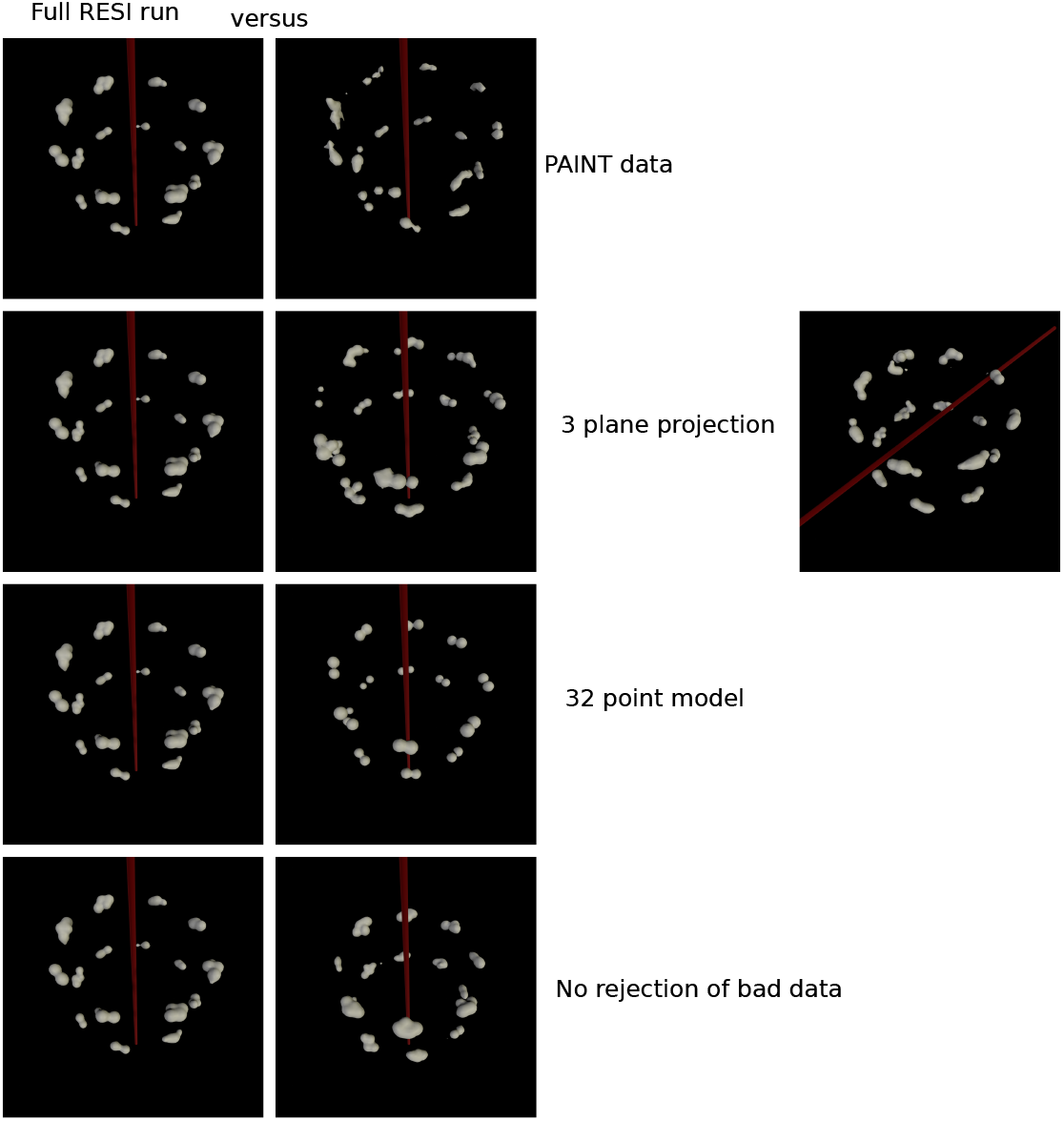
Comparing performance of SQUASSH on the RESI data. Left hand column shows the RESI data analysed with SQUASSH as shown in the main paper. In the right hand column the first row shows the result on the raw PAINT data without RESI processing, the second row shows the results with three plane projection instead of six plane rendering. With three plane projection, in about 30% of the runs, SQUASSH does not get the optimal stretch axis, with a sample shown in the rightmost panel. In all cases the axis is orthogonal to the correct direction. The third row shows the result with 32 points rather than 700 in the structural model, and the fourth row shows the result with no auxiliary loss, i.e. when the algorithm is not able to reject data to improve performance.

**Fig. 4.**
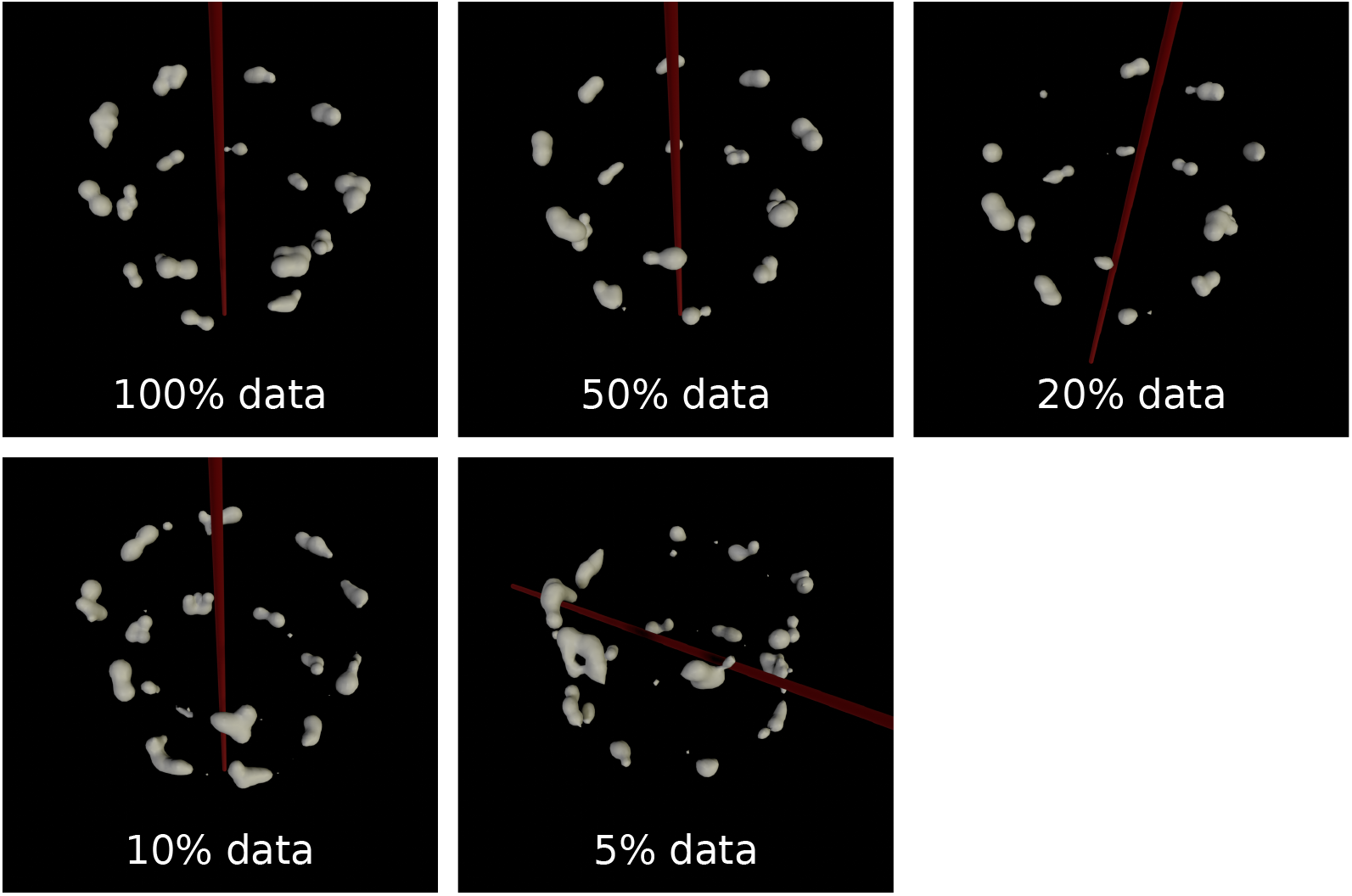
Comparing performance of SQUASSH with amount of data provided to the algorithm. For the RESI data, while performance is substantially unaffected down to 50% of the data, performance then gradually degrades. However the general features of the structural model (two rings and eightfold symmetry) can be observed down to 5% of the data (61 images), indicating that for relatively simple structures reasonable quality reconstructions can be achieved with under 100 images.

**Fig. 5.**
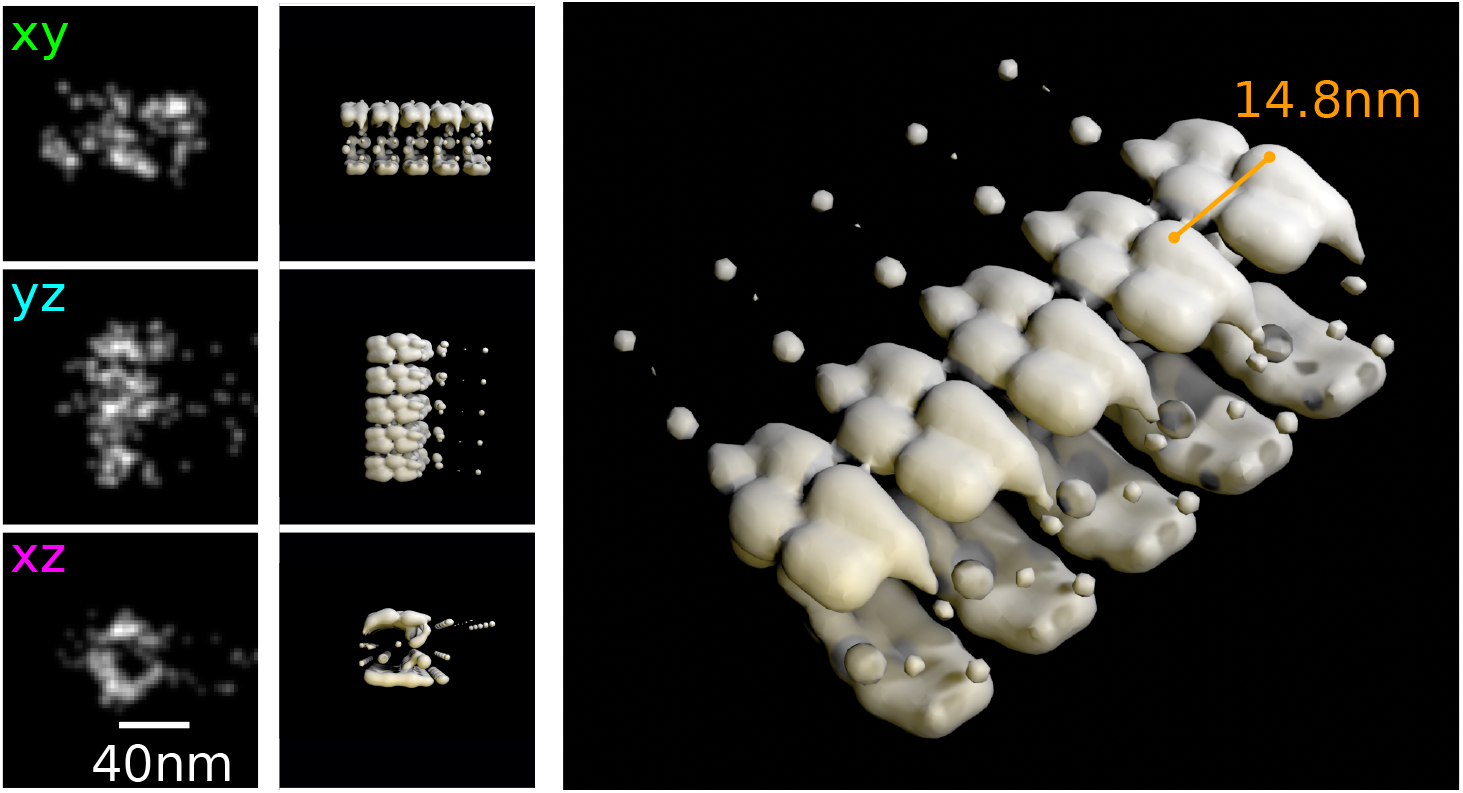
SQUASSH analysis of microtubule data finds a repeat distance of 7.4 nm when run with a smaller patch size, illustrating how the measured local repeat distance can be biased by the random scatter present in SMLM data. The parameters for the microtubule run are as described in Sec. 4.3.1, but with a segment length of 64 nm.

**Fig. 6.**
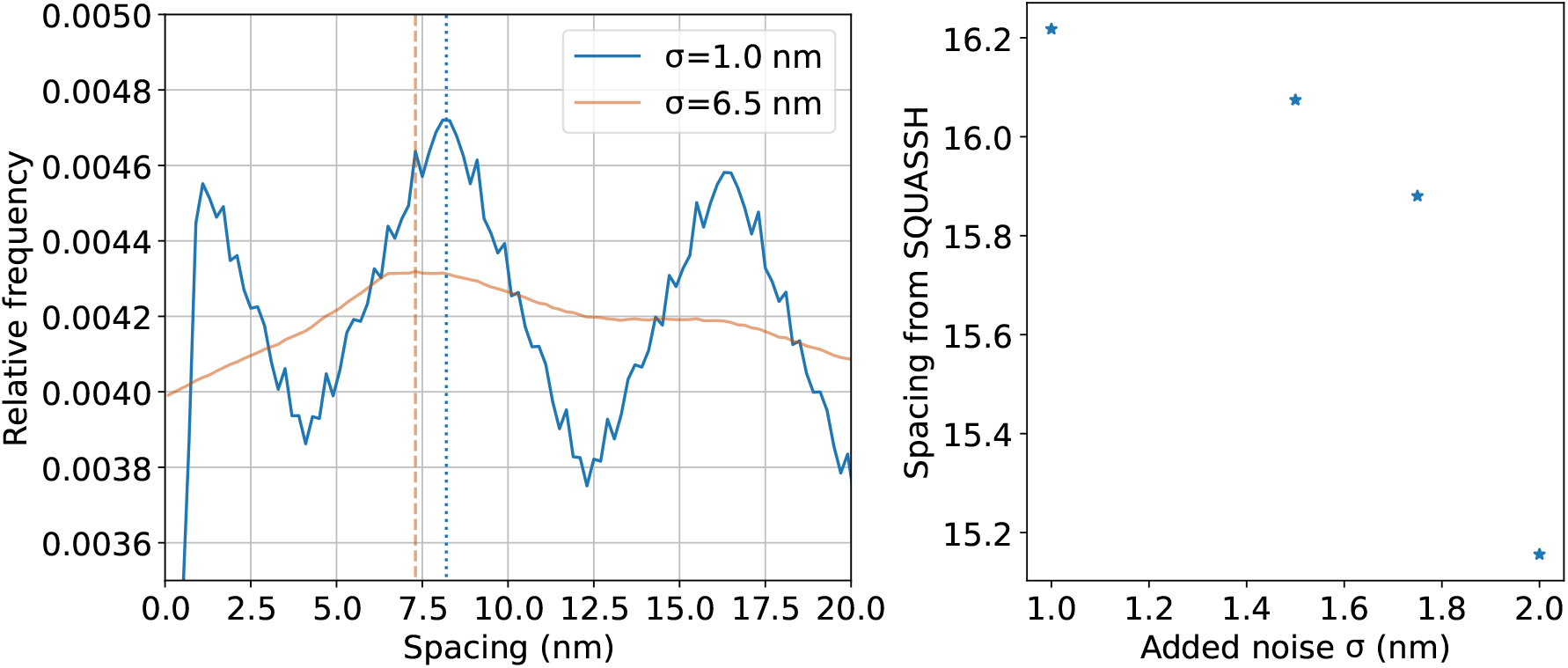
Repeat distance of a periodic structure is affected by the scatter in the results. Left: the 3D structure of a microtubule was simulated with different levels of noise added to the positions, and a histogram of distances between every pair of points was computed. With a small amount of added noise, there is a clear peak at 8.2nm (indicated with the vertical line), corresponding to the repetition distance of the structure. With a large amount of added noise, the mode of the distances shifts to a slightly smaller value (around 7.3nm). Right: SQUASSH was applied to the simulated microtubules and the graph shows repeat distance of SQUASSH repeating unit versus sigma of Gaussian scatter applied to simulated microtubule structure (400 patches were simulated and fitted).

**Fig. 7.**
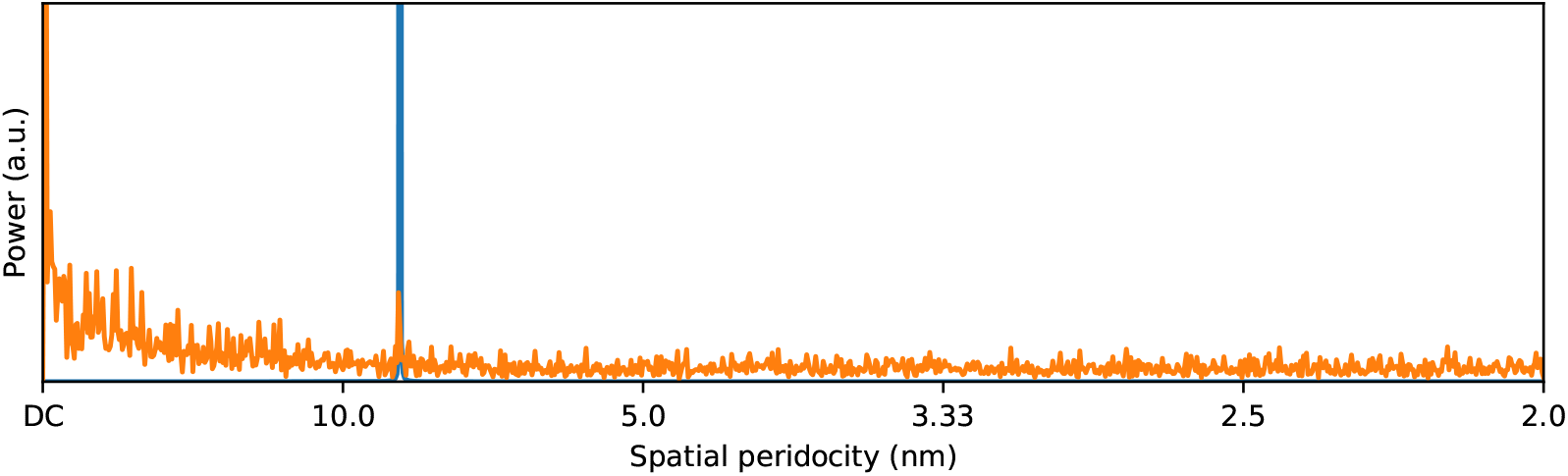
Fourier analysis of microtubules shows a significant frequency component corresponding to 8.4 nm periodicity. For this analysis, 6 straight segments of microtubles 2000 nm in length were projected on to the long axis and then rendered. The power spectra of the segments were computed and then averaged. There is a significant peak at 8.4 nm, highlighted by the vertical blue line.

**Fig. 8.**
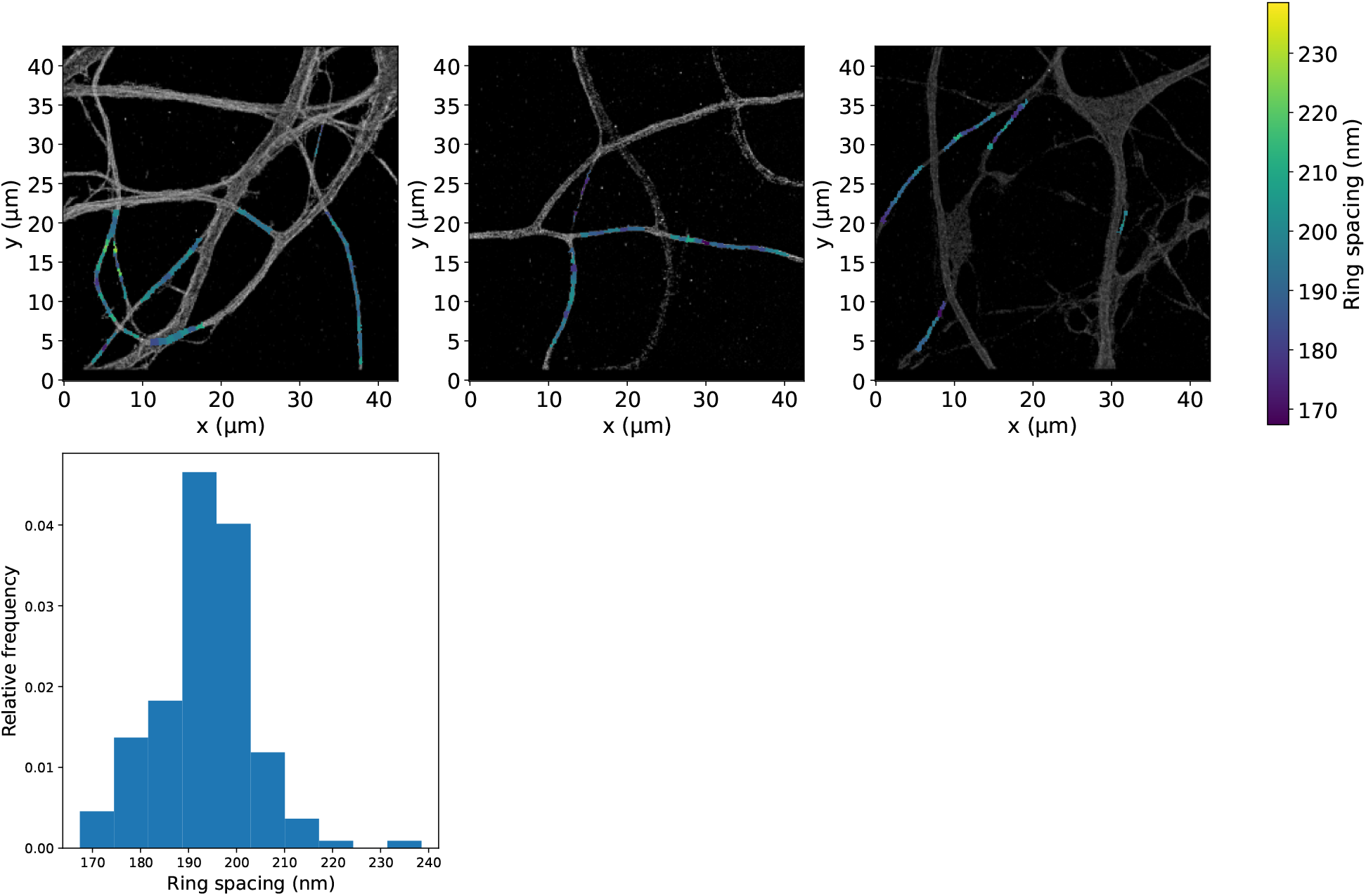
Spacing between spectrin rings in axons found using SQUASSH. Top row: reconstructed SMLM images of axons with SQUASSH analysis. Areas to analyse were selected based on them having a single ring structure rather than multiple joined together, or crossing. Spacing values are superimposed in colour, illustrating the variation in ring spacing across the sample. Bottom row: histogram of ring spacings, with a mean value of 94 nm.

**Fig. 9.**
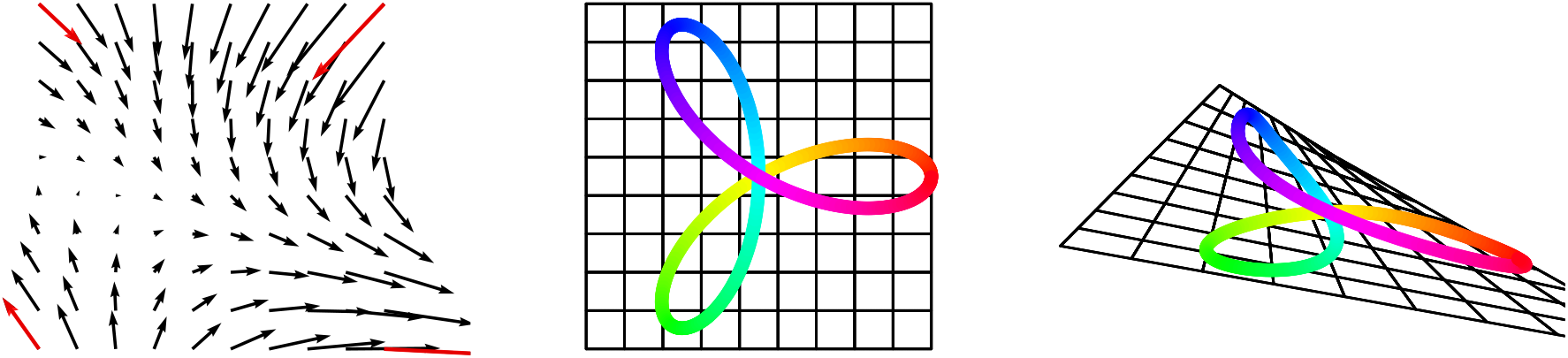
Illustration of the general distortion model in 2D. Left: the distortion is parameterised by a vector at each corner of a square (shown in red), so the 2D parameterisation has 8 parameters. Vectors at an arbitrary positions (black) are found by independent bilinear interpolation of the x and y values of the vectors. Centre: an example shape and grid. Right: all points in the shape and grid are distorted by adding the distortion vectors from the model. Note how the distortion model allows for lengthening and shortening of the shape lobes as well as changing the angles between the lobes.

**Fig. 10.**
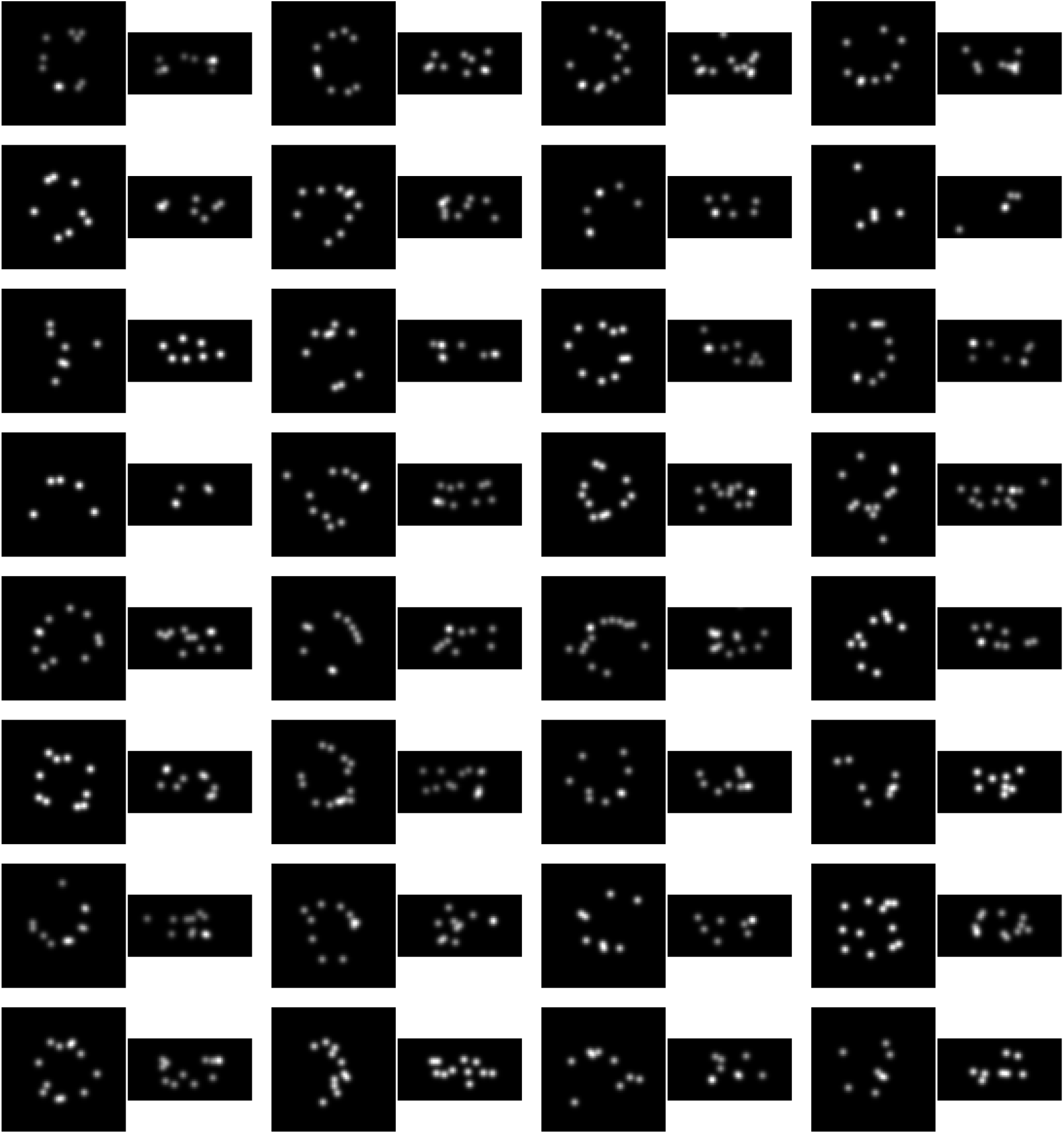
Examples of nuclear pore complexes from ^1^ processed with RESI, which were identified as invalid by SQUASSH. Nuclear pore complexes are shown with XY and XZ projections. In all of the examples, significant problems can be seen, such as missing sections of the ring, the upper or lower ring missing, severe distortion or segmentation errors giving spurious fluorophores.

**Fig. 11.**
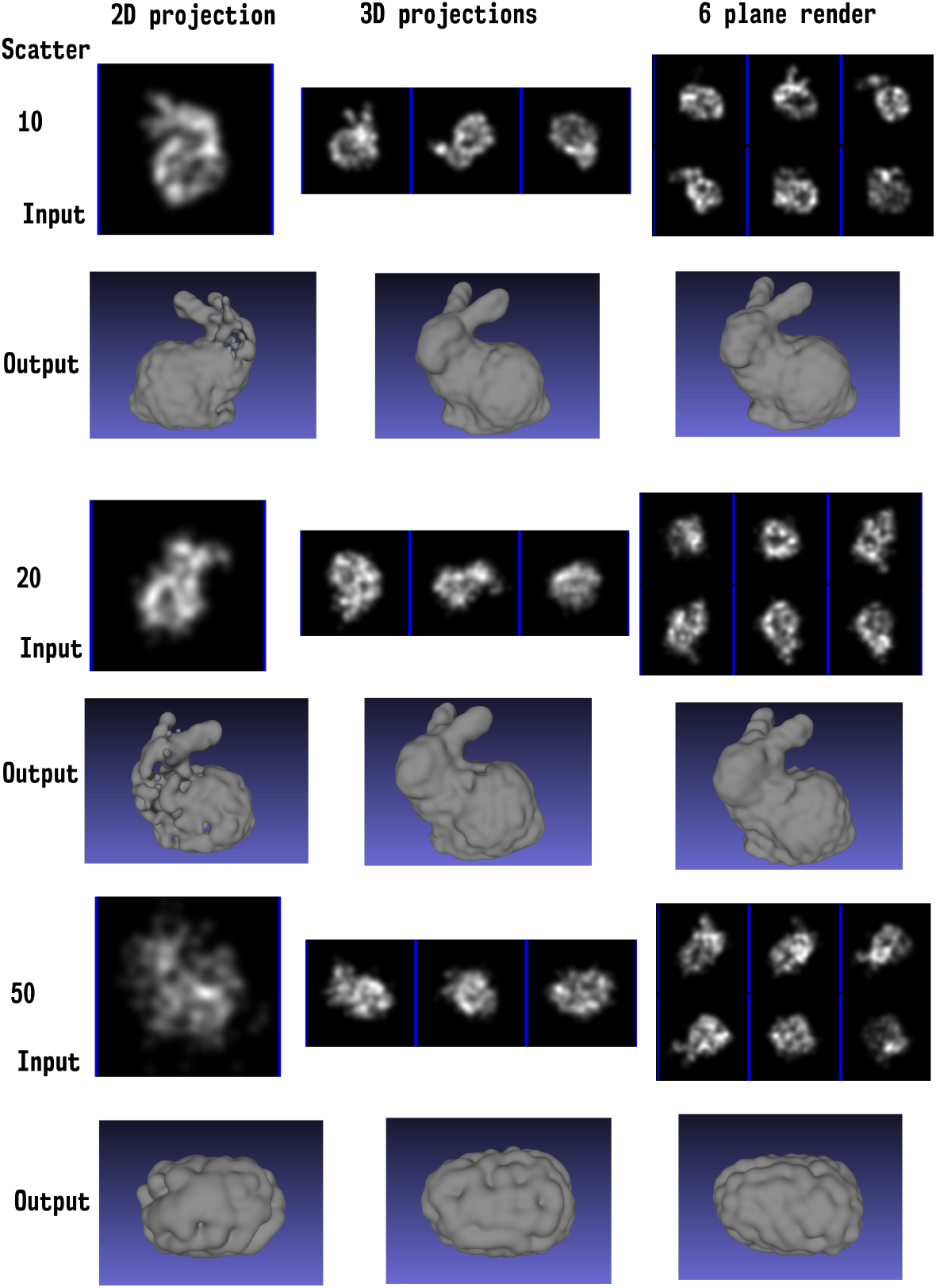
Impact of the scatter of points and the rendering approach on the output structural model. Columns group different rendering approaches, and rows 10, 20 and 50 pixel sigma scatter. For each combination an example of a training input image is shown in the Input row, and the final optimised structural model is shown in the Output row. Performance of SQUASSH decreases with increased scatter in the input data, and increases as the projection method becomes more informative, with a single 2D projection being least informative, three 3D projections being intermediate and a 6 plane rendering which encodes z information in the intensity being the best performer.

**Fig. 12.**
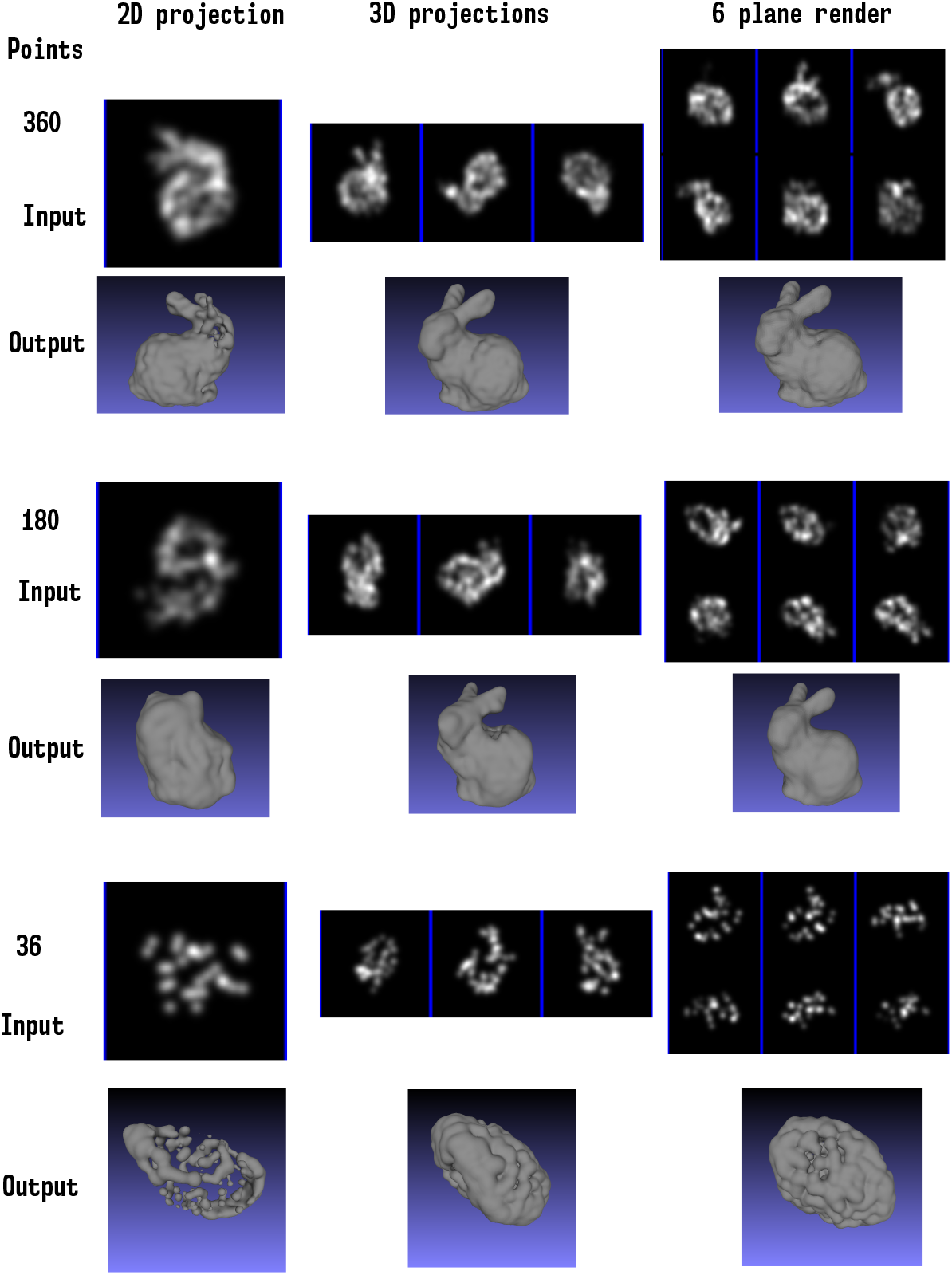
Impact of labelling efficiency and rendering approach on the output structural model. Columns group different rendering approaches, and rows the number of points in the structural model, with the top being 360, the middle being 180 and the bottom being 36. For each combination an example of a training input image is shown in the Input row, and the final optimised structural model is shown in the Output row. Performance of SQUASSH decreases when there are fewer points in the input model, though with a highly informative rendering the quality of the output model can be retained even when the labelling efficiency has dropped substantially (for the six plane render, there is little degradation in quality dropping from 360 to 180 points, while for the 2D projections the quality of the output drops substantially).

#### 1 Movie caption

Demonstration of how the rotational fitting is optimised alongside the structural model, and learning of the underlying manifold. SQUASSH was applied to simulated data derived from the Stanford bunny. For each element of data, fluorophore positions were derived from the mesh vertices with scatter and dropout and then a random rotation and translation was applied. The optimisation was stopped at three different stages, the rotation network was extracted and its performance was tested by inputting a new data element while smoothly varying the rotation. Rows show different stages of the optimisation (top row being earliest, bottom row being latest). From left to right, the first column shows data data elements being inputted. The second column shows the input projection (using the blur at that stage of the optimisation), the third column shows the ground truth structural model at the pose given by the rotation network, and the fourth column shows the current structural model at the pose given by the rotation network. This demonstrates the underlying rotation manifold being learned: as the input data is rotated, the output rendering generally rotates in a similar way. The network can predict many rotations correctly at the beginning, but at some positions (especially where there is close to a reflective ambiguity, the results are not correct. However, by the bottom row correct rotations are predicted for all poses.

#### 2 Movie caption

Optimisation of a structure using SQUASSH analysis. The fluorophores are initiated to random positions and then optimise over time. Left: each fluorophore is rendered as a sphere. Right: each fluorophore is rendered as a Gaussian, and a marching cubes algorithm is used to render an isosurface.

#### 3 Movie caption

Optimisation of the structure of nup96 within the nuclear pore complex. The structure is initiated with 35 fluorophores at random positions and then optimised. The postions of each fluorophore are then re-seeded with a total of 700 fluorophores for the whole image, and the optimisation proceeds. This process allows us to create a probability density map for fluorophore positions within nup96 that can be intuitively represented as an isosurface, without requiring knowledge of the precise number of fluorophores within a given structure.

#### 4 Movie caption

Structure of spectrin rings within axons. Three repeat units are shown, with a random sampling of points within the pca being made and then smoothly interpolating between them. Note that the quality at the top and bottom of the structure is degraded compared to the sides, as the structure is being observed in the same axial orientation every time, and therefore the optimisation cannot benefit from observations at different angles. It can be seen that the shape and size of the rings vary considerably.

#### 5 Movie caption

In telophase the chromosomes arrive at the cell poles and a new nuclear membrane forms around each of the sets of chromosomes. Lattice light sheet microscopy data is modelling as arising from two structures separated by a variable distance along an axis, with one structure also able to rotate relative to that axis. This allows fixed cell observations to be reordered to create a virtual dividing cell, and the distance between the two chromatin disks in each observation to be identified.

