## Supplementary note and figures for "Extracting biological structure and heterogeneity from the nano to the macro scale"

**This PDF file includes:**

Figures S1 to S12
Captions for Movies S1 to S5

**Other Supplementary Materials for this manuscript:**

Movies S1 to S5

### Supplementary Note

#### 1 Reflection/rotation ambiguity

We reconstruct 3D shapes by predicting Euclidean transformations for a 3D structural model. If the reconstruction uses only one projection for each predicted pose, then the shape can only be determined up to a reflection. Proof:

Consider an abitrary point $\mathbf{X} = [\alpha X, \alpha Y, \alpha Z, \alpha]$, $\mathbf{X} \in \mathbb{P}^3$. For simplicity but without loss of generality we will consider only $\alpha = 1$. The point is transformed by and a Euclidean transform $\mathbf{E} : \mathbb{P}^3 \longrightarrow \mathbb{P}^3$:

$$\mathbf{E} = \left[\begin{array}{c|c} \mathbf{R} & \mathbf{T} \\ \hline \mathbf{0} & \mathbf{1} \end{array}\right],$$

where $\mathbf{R}$ and $\mathbf{T}$ are a rotation and translation in $\mathbb{R}^3$. With affine projection (how microscopes work), the 2D position, $\mathbf{u}$, is:

$$\mathbf{u} = \underbrace{\begin{bmatrix} 1 & 0 & 0 \\ 0 & 1 & 0 \end{bmatrix}}_{\substack{\Pi:\mathbb{P}^2 \rightarrow \mathbb{R}^2 \\ \text{Projection for } \alpha=1}} \underbrace{\begin{bmatrix} k_1 & k_2 & k_3 \\ 0 & k_4 & k_5 \\ 0 & 0 & 1 \end{bmatrix}}_{\substack{\mathbf{k}:\mathbb{P}^2 \rightarrow \mathbb{P}^2 \\ \text{Camera}}} \underbrace{\begin{bmatrix} 1 & 0 & 0 & 0 \\ 0 & 1 & 0 & 0 \\ 0 & 0 & 0 & 1 \end{bmatrix}}_{\substack{\mathbf{P}:\mathbb{P}^3 \rightarrow \mathbb{P}^2 \\ \text{Affine projection}}} \mathbf{EX},$$

where $\mathbf{k}$ is an arbitrary camera calibration matrix. Composing the projections and camera yielding $\mathbf{K} = \Pi \mathbf{k} \mathbf{P}$ gives:

$$\mathbf{u} = \underbrace{\begin{bmatrix} k_1 & k_2 & 0 & k_3 \\ 0 & k_4 & 0 & k_5 \end{bmatrix}}_{\mathbf{K}} \left[\begin{array}{c|c} \mathbf{R} & \mathbf{T} \\ \hline \mathbf{0} & \mathbf{1} \end{array}\right] \mathbf{X}.$$

Defining $\mathbf{X}' = [-X, -Y, -Z, 1]$ as the reflection of X. then it is clear that:

$$\mathbf{u} = \begin{bmatrix} k_1 & k_2 & 0 & k_3 \\ 0 & k_4 & 0 & k_5 \end{bmatrix} \left[\begin{array}{c|c} -\mathbf{R} & \mathbf{T} \\ \hline \mathbf{0} & \mathbf{1} \end{array}\right] \mathbf{X}'.$$

That is not a valid Eucildean transform, bit since $\mathbf{K}$ has a zero third column, the result is invariant to changes in the third row of the rotation matrix, therefore we can pick $\mathbf{R}' = \begin{bmatrix} -1 & 0 & 0 \\ 0 & -1 & 0 \\ 0 & 0 & 1 \end{bmatrix} \mathbf{R}$, which also gives:

$$\mathbf{u} = \begin{bmatrix} k_1 & k_2 & 0 & k_3 \\ 0 & k_4 & 0 & k_5 \end{bmatrix} \left[\begin{array}{c|c} \mathbf{R}' & \mathbf{T} \\ \hline \mathbf{0} & \mathbf{1} \end{array}\right] \mathbf{X}'.$$

Therefore if we mirror the structure and rotate the pose by 180° around the Z azis, the projected image will be identical.

It is important to note that this holds only because the third columns of $\mathbf{k}$ is zero, allowing us to substitute a rotation matrix which mirrors two of the axes. This does not hold if there are projections along multiple different axes.

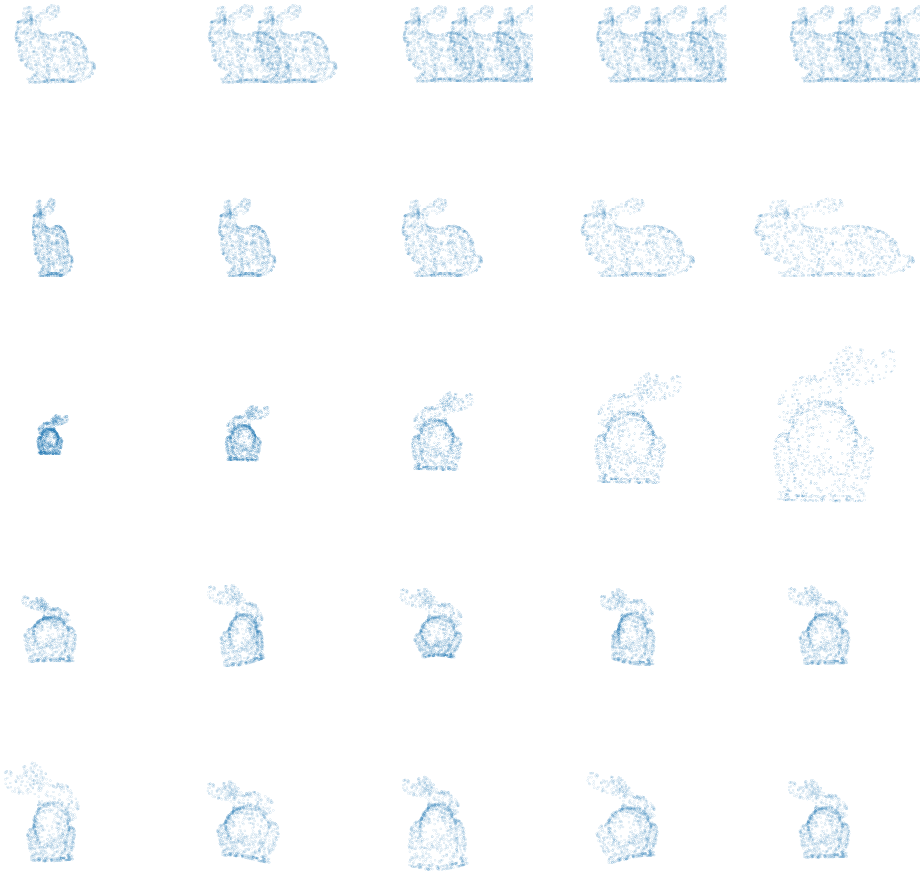

**Fig. 1** Illustration of different heterogeneity parameterisations. Following Fig. 1 we use the Stanford Bunny as an exemplar structural model, with the primary axis aligned with x. Row 1: variable duplication along an axis. Row 2: elongation along an exis. Row 3: expansion orthongonal to an axis. Rows 4 and 5: various different angle dependent scaling orthogonal to an axis.

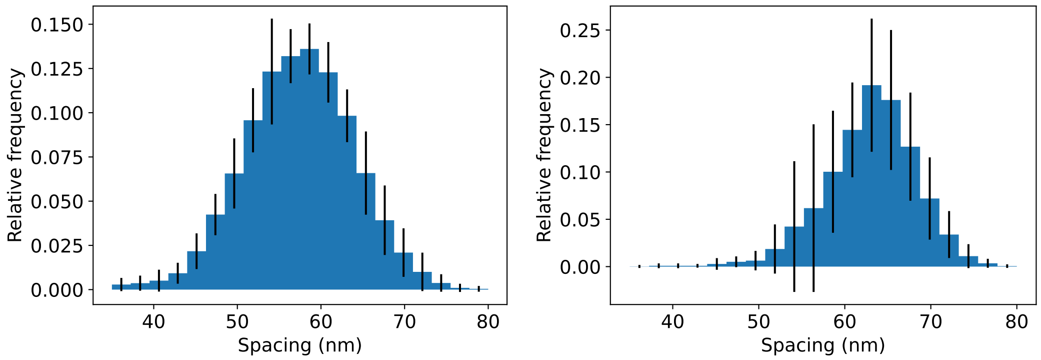


**Fig. 2** Mean histograms of nuclear pore complexsx ring spacings from the RESI dataset (left) and the 4Pi-STORM dataset (right). SQUASSH was rerun 10 times, in order to compute the mean value of histogram bins along with error bars as ±1 standard deviation.

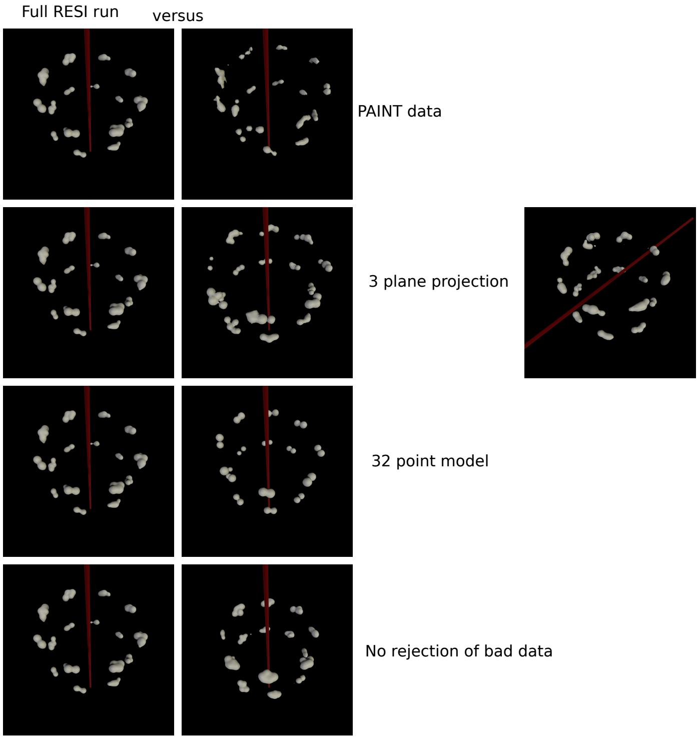


**Fig. 3** Comparing performance of SQUASSH on the RESI data. Left hand column shows the RESI data analysed with SQUASSH as shown in the main paper. In the right hand column the first row shows the result on the raw PAINT data without RESI processing, the second row shows the results with three plane projection instead of six plane rendering. With three plane projection, in about 30% of the runs, SQUASSH does not get the optimal stretch axis, with a sample shown in the rightmost panel. In all cases the axis is orthogonal to the correct direction. The third row shows the result with 32 points rather than 700 in the structural model, and the fourth row shows the result with no auxiliary loss, i.e. when the algorithm is not able to reject data to improve performance.

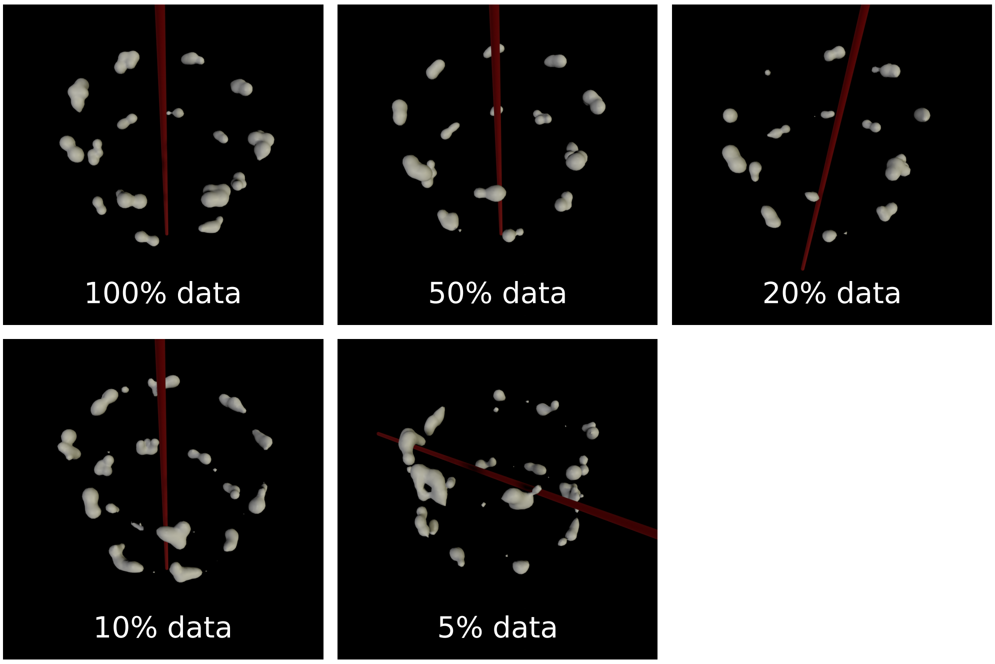


**Fig. 4** Comparing performance of SQUASSH with amount of data provided to the algorithm. For the RESI data, while performance is substantially unaffected down to 50% of the data, performance then gradually degrades. However the general features of the structural model (two rings and eightfold symmetry) can be observed down to 5% of the data (61 images), indicating that for relatively simple structures reasonable quality reconstructions can be achieved with under 100 images.

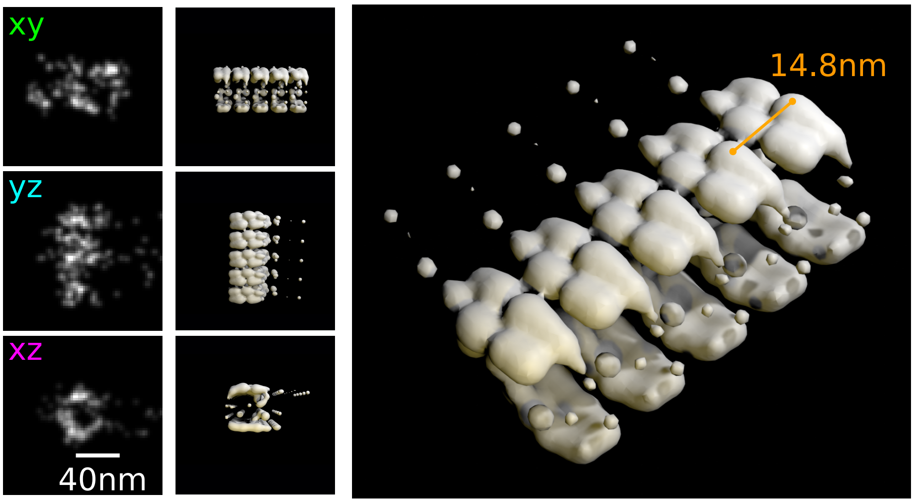


**Fig. 5** SQUASSH analysis of microtubule data finds a repeat distance of 7.4 nm when run with a smaller patch size, illustrating how the measured local repeat distance can be biased by the random scatter present in SMLM data. The parameters for the microtubule run are as described in Sec. 4.3.1, but with a segment length of 64 nm.

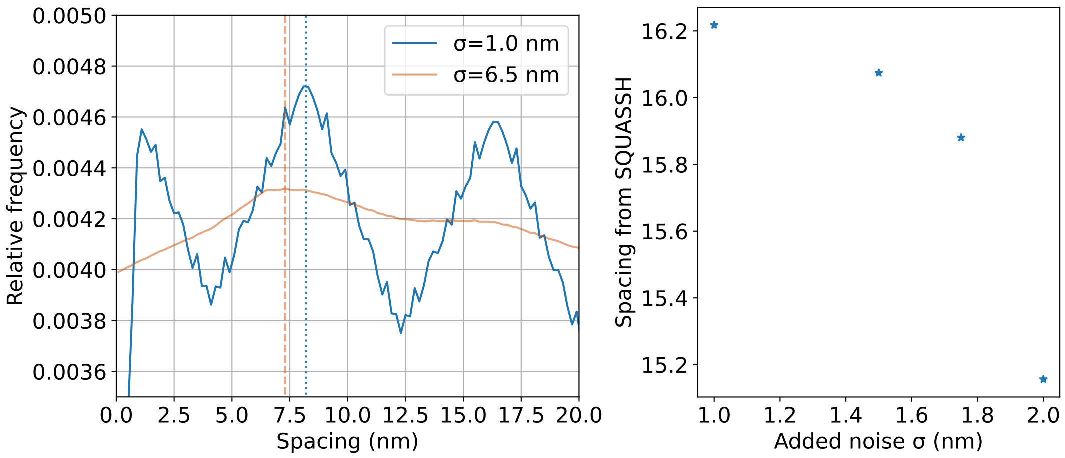


**Fig. 6** Repeat distance of a periodic structure is affected by the scatter in the results. Left: the 3D structure of a microtubule was simulated with different levels of noise added to the positions, and a histogram of distances between every pair of points was computed. With a small amount of added noise, there is a clear peak at 8.2nm (indicated with the vertical line), corresponding to the repetition distance of the structure. With a large amount of added noise, the mode of the distances shifts to a slightly smaller value (around 7.3nm). Right: SQUASSH was applied to the simulated microtubules and the graph shows repeat distance of SQUASSH repeating unit versus sigma of Gaussian scatter applied to simulated microtubule structure (400 patches were simulated and fitted).

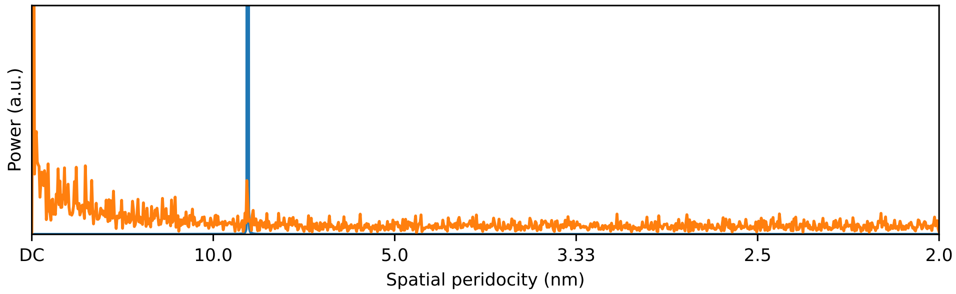


**Fig. 7** Fourier analysis of microtubules shows a significant frequency component corresponding to 8.4 nm periodicity. For this analysis, 6 straight segments of microtubles 2000 nm in length were projected on to the long axis and then rendered. The power spectra of the segments were computed and then averaged. There is a significant peak at 8.4 nm, highlighted by the vertical blue line.

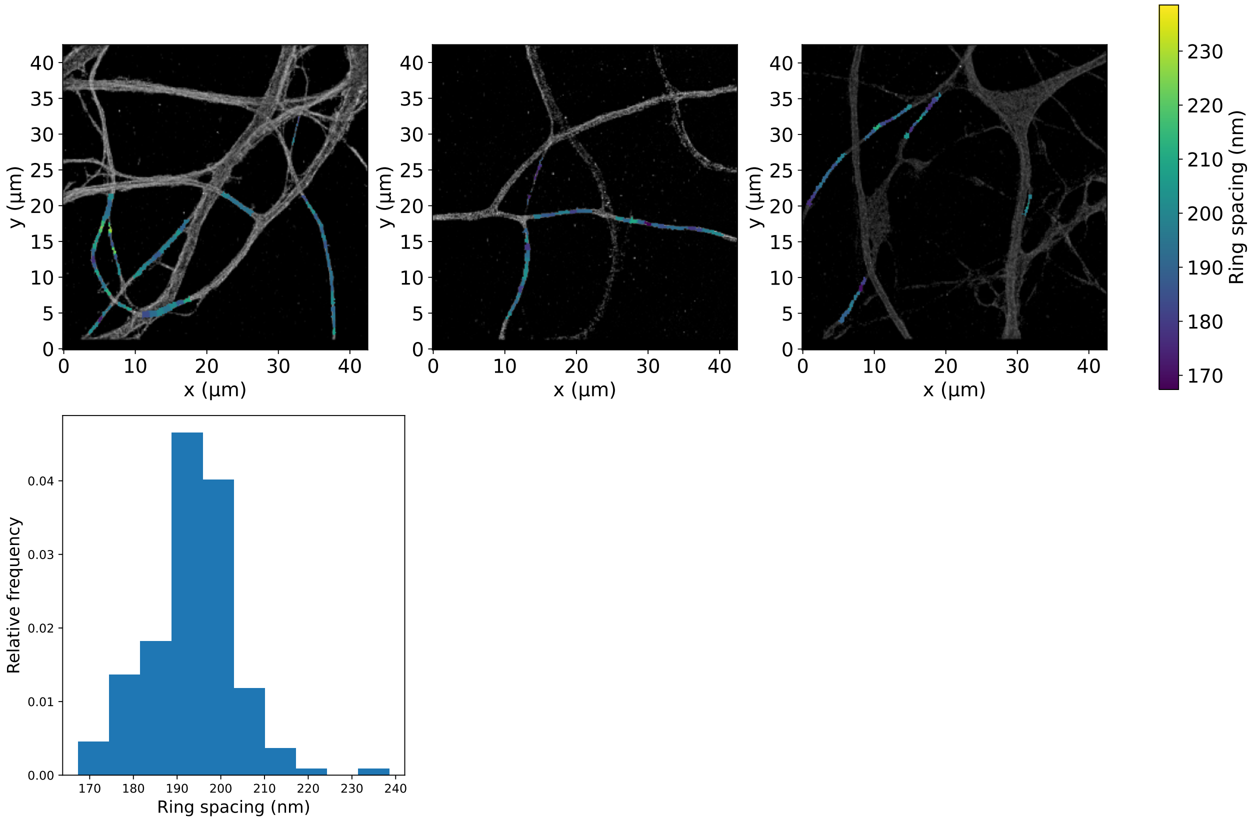


**Fig. 8** Spacing between spectrin rings in axons found using SQUASSH. Top row: reconstructed SMLM images of axons with SQUASSH analysis. Areas to analyse were selected based on them having a single ring structure rather than multiple joined together, or crossing. Spacing values are superimposed in colour, illustrating the variation in ring spacing across the sample. Bottom row: histogram of ring spacings, with a mean value of 94 nm.

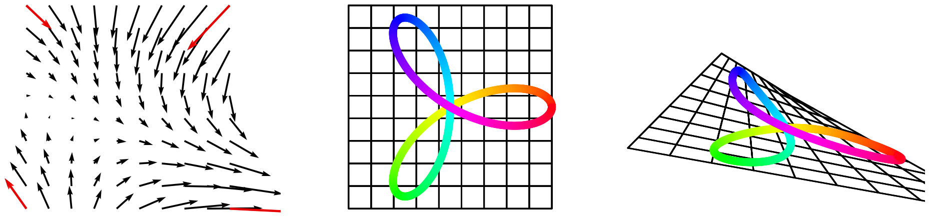

**Fig. 9** Illustration of the general distortion model in 2D. Left: the distortion is parameterised by a vector at each corner of a square (shown in red), so the 2D parameterisation has 8 parameters. Vectors at an arbitrary positions (black) are found by independent bilinear interpolation of the x and y values of the vectors. Centre: an example shape and grid. Right: all points in the shape and grid are distorted by adding the distortion vectors from the model. Note how the distortion model allows for lengthening and shortening of the shape lobes as well as changing the angles between the lobes.

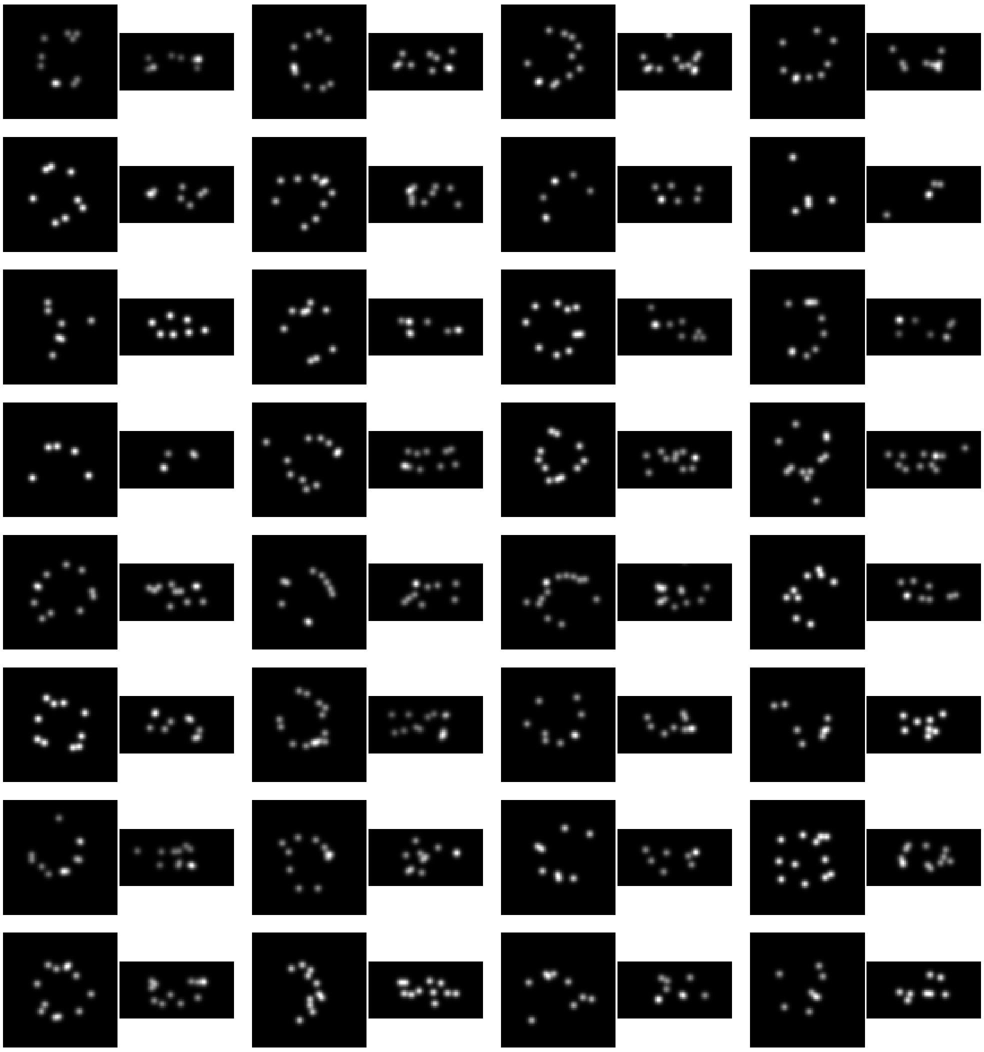

**Fig. 10** Examples of nuclear pore complexes from[1] processed with RESI, which were identified as invalid by SQUASSH. Nuclear pore complexes are shown with XY and XZ projections. In all of the examples, significant problems can be seen, such as missing sections of the ring, the upper or lower ring missing, severe distortion or segmentation errors giving spurious fluorophores.

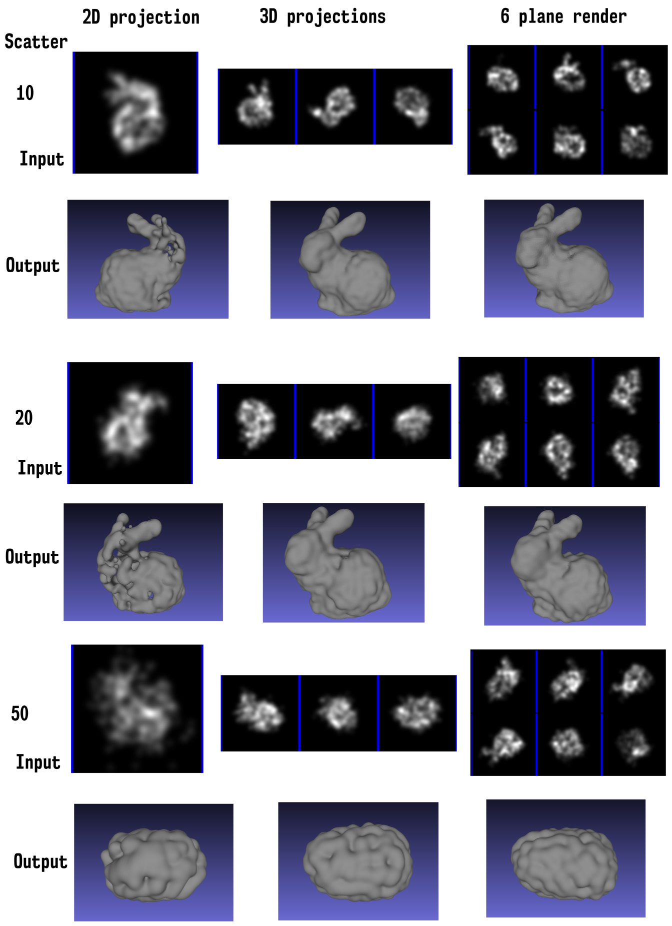


**Fig. 11** Impact of the scatter of points and the rendering approach on the output structural model. Columns group different rendering approaches, and rows 10, 20 and 50 pixel sigma scatter. For each combination an example of a training input image is shown in the Input row, and the final optimised structural model is shown in the Output row. Performance of SQUASSH decreases with increased scatter in the input data, and increases as the projection method becomes more informative, with a single 2D projection being least informative, three 3D projections being intermediate and a 6 plane rendering which encodes z information in the intensity being the best performer.

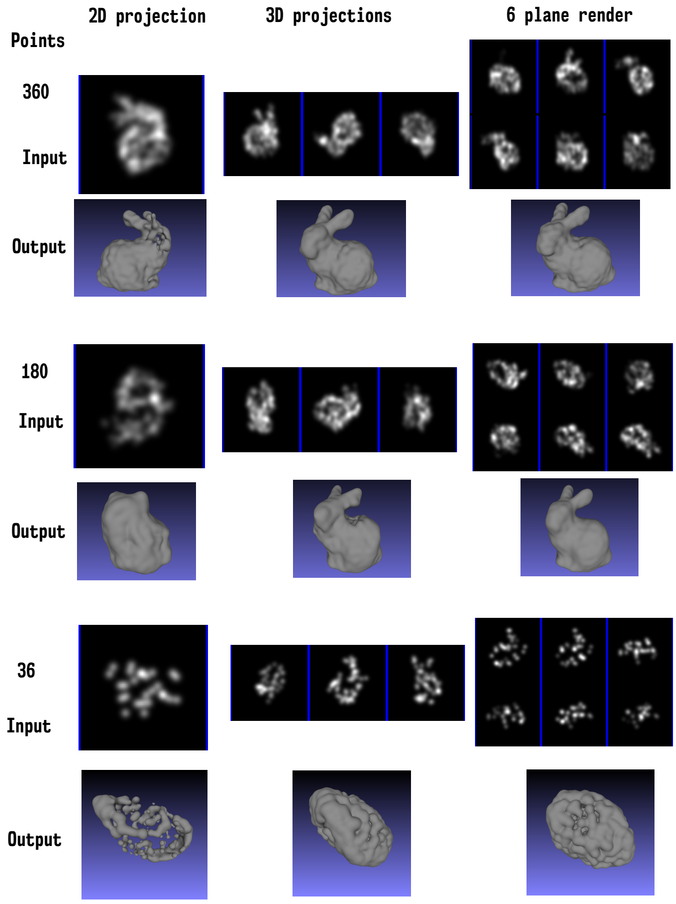


**Fig. 12** Impact of labelling efficiency and rendering approach on the output structural model. Columns group different rendering approaches, and rows the number of points in the structural model, with the top being 360, the middle being 180 and the bottom being 36. For each combination an example of a training input image is shown in the Input row, and the final optimised structural model is shown in the Output row. Performance of SQUASSH decreases when there are fewer points in the input model, though with a highly informative rendering the quality of the output model can be retained even when the labelling efficiency has dropped substantially (for the six plane render, there is little degradation in quality dropping from 360 to 180 points, while for the 2D projections the quality of the output drops substantially).
